# Cell-cycle dependence of bursty gene expression: insights from fitting mechanistic models to single-cell RNA-seq data

**DOI:** 10.1101/2024.01.10.574820

**Authors:** Augustinas Sukys, Ramon Grima

## Abstract

Bursty gene expression is characterised by two intuitive parameters, burst frequency and burst size, the cell-cycle dependence of which has not been extensively profiled at the transcriptome level. In this study, we estimate the burst parameters per allele in the G1 and G2/M cell-cycle phases for thousands of mouse genes by fitting mechanistic models of gene expression to mRNA count data, obtained by sequencing of single cells whose cell-cycle position has been inferred using a deep-learning method. We find that upon DNA replication, the median burst frequency approximately halves, while the burst size remains mostly unchanged. Genome-wide distributions of the burst parameter ratios between the G2/M and G1 phases are wide, indicating substantial heterogeneity in transcriptional regulation patterns. We also observe a significant negative correlation between the burst frequency and size ratios, suggesting that regulatory processes do not independently control the burst parameters. Finally, we argue that to accurately estimate the burst parameter ratios, mechanistic models must explicitly account for gene copy number variation and extrinsic noise due to the coupling of transcription to cell age within the cell cycle, but corrections for technical noise due to imperfect capture of RNA molecules in sequencing experiments are less critical.

## Introduction

Gene expression occurs in random bursts of transcription, associated with alternating active and inactive promoter states [1, 2]. By tagging mRNA molecules using fluorescent reporters, live-cell imaging reveals a train of pulses of fluorescent intensity whose mean height is proportional to the mean number of transcripts produced when the gene is actively transcribing (the mean burst size) and whose frequency reflects how often transcription occurs (the burst frequency) [3]. These two burst parameters offer a simple, intuitive and practical quantitative description of gene expression and hence their estimation has been the focus of many studies (for reviews see [4–6]).

While live-cell measurements are ideal to understand transcription, they are challenging because they are low-throughput and require genome editing [7, 8]. To overcome this challenge, one can leverage the inherent heterogeneity generated by bursting within a population of cells. Acquiring a distribution of mRNA counts per cell is nowadays relatively straightforward through techniques such as single-molecule fluorescence in situ hybridization (smFISH) and single-cell sequencing. By fitting these measured distributions to the distributions predicted by mechanistic models of gene expression (where the rate parameters determine the speed of molecular processes) one can then infer the burst size and frequency. Common examples of such models are the two-state telegraph model [1, 9] which predicts both unimodal and bimodal mRNA count distributions, and the simpler bursty model that predicts unimodal negative binomial distributions [10].

The vast majority of studies reporting estimates of the burst size and frequency have used smFISH data [11– 14] because it is widely considered to be the gold standard for the accurate measurement of the number of RNA molecules for a specific gene in single cells. However, many of these studies suffer from a major limitation: *they implicitly assume that the burst parameters are the same for each cell in the sample*. Equivalently, it is assumed that the observed inter-cell differences in the transcript numbers are generated by the uncertainty in the timing of biochemical events (intrinsic noise) [15, 16]. However, generally this is not the case — burst parameters vary from one cell to another because of differences in the number of cellular components such as RNA polymerases, transcription factors and other molecules playing key roles in transcription. These parameter fluctuations are commonly referred to as extrinsic noise, since they arise independently of a gene of interest but act on it. Thus, the differences in transcript numbers between cells are due to both intrinsic and extrinsic noise, although the latter has been suggested to be the dominant source of heterogeneity [17, 18].

One might suppose that the burst parameters estimated using models that assume perfectly identical cells can be approximately interpreted as the parameter values averaged over all cells. However, a recent study conclusively showed this is not the case. In Ref. [19] it was shown that there exist systematic biases in parameters inferred using the telegraph model when its distribution or moments are fitted to those from simulated single-allele mRNA data from a population of cells with extrinsic noise in one of the parameters. Whether the burst parameters are over- or underestimated was found to depend on the source of extrinsic noise (which parameter is the most variable amongst cells) and the mode of transcriptional activity. In reality, these systematic biases are likely to be larger when non-allele-specific mRNA count data is used for inference because the cell-to-cell differences in the gene copy numbers (which occur when cells are in different cell-cycle phases) can be a strong determinant of gene expression noise [20].

Together, these studies show that it is difficult to obtain a reliable quantitative picture of gene expression by fitting models to single-cell data unless the inference procedure appropriately corrects for the changes of gene copy number with the cell-cycle phase and for extrinsic noise due to other exogenous factors that vary within each phase. A number of smFISH-based studies have suggested that the burst frequency is principally affected by the cell-cycle phase [20–22] while the burst size scales with the cell volume [21, 23], which leads to concentration homeostasis [24]. Note that cell volume varies considerably within a cell-cycle phase and hence it is possibly a dominant source of extrinsic noise within each phase; other extrinsic noise sources include mitochondrial variability [25] and Ca2+ signalling [26]. However, even if sophisticated modelling frameworks that include all the principal sources of extrinsic noise can be developed and used to fit the mRNA count distributions measured using smFISH, this would not lead to a transcriptome-wide quantitative understanding of gene expression as such data is typically limited to a relatively small number of genes of interest.

A possible way of circumventing this problem involves the use of data generated by single-cell RNA sequencing (scRNA-seq). This allows the comprehensive profiling of large numbers of cells on a genome-wide level, thus providing an unprecedented insight into cell-to-cell variability [27–29]. While it has primarily been used to distinguish between different cell types in a sample [30–33], to provide insights into the dynamics of cell state switching [34–36] and to reconstruct gene-regulatory networks [37–41], in recent years it has also been used to study the inherent stochasticity of gene expression [19, 42–48]. Despite its potential for transcriptome-wide inference, scRNA-seq data has one major disadvantage compared to smFISH data, namely the high amount of technical (non-biological) noise [49, 50]. In droplet-based sequencing this can arise from the imperfect capture of all RNA molecules (especially from lowly expressed genes), amplification errors and non-uniquely identifiable reads. Although technical noise is reduced by the use of data that incorporates unique molecular identifiers (UMIs) [51, 52] — random nucleotide sequences that tag an individual mRNA molecule and thus enable separation between the original molecules and amplification duplicates derived from the cDNA library amplification [53–55] — this is not a panacea. A comparison of the probability distributions of mRNA counts for five genes measured using both smFISH and a UMI-based scRNA-seq platform implies there is a non-negligible fraction of transcripts in each cell that are not captured by the sequencing technology (Extended Figure 7 of [43]). Considering the limitations of inference already identified using smFISH data, it is clear that the deduction of quantitative models of bursty gene expression using scRNA-seq data would need an inference procedure that corrects for the cell-cycle phase, extrinsic noise within each phase and technical noise. This trifarious correction is missing from the literature. Current methods of inferring burst parameters from scRNA-seq data either ignore/cannot properly account for extrinsic and technical noise [42, 43] or correct only for extrinsic noise [19], or correct for certain types of extrinsic noise and technical noise but not for the variation in gene copy number over the cell cycle [44, 45, 48, 56].

In this paper, we develop an inference framework that takes into account the cell-cycle phase, extrinsic noise due to factors varying within each phase and technical noise, and use it to investigate how the burst frequency and size are modulated over the cell cycle in eukaryotic genes. While previous studies sought to answer this question using smFISH-based approaches for a few tens of genes, we use previously published scRNA-seq data [57] to understand the cell-cycle dependence of the bursty expression of about 1000 mouse genes. The overall approach is illustrated in Figure 1. Sequencing data provides spliced and unspliced read counts for over 10^4^ genes in undifferentiated mouse embryonic stem cells. DeepCycle [57] uses a deep-learning approach to map the cycling patterns in the unspliced-spliced RNA space of cell-cycle-related genes to a parameter *θ* for each cell, which gives the cell age: *θ* = 0 means a newborn cell, while *θ* = 1 indicates a cell that is about to divide via mitosis (hence *θ* is a rough proxy for cell size). From this information, it is also possible to determine the cell-cycle phase which agrees with the phases according to the DNA content given by Fluorescence-Activated Cell Sorting (FACS). Given this joint information of mRNA counts, cell age *θ* and cell-cycle phase for each cell, we fit two classes of mechanistic models of stochastic transcription (age-independent and age-dependent models) to the data, from which we estimate the burst parameters. We show that the most robust inference strategy involves using age-dependent models to estimate the ratios of the burst parameters between the G1 and G2/M phases, rather than their phase-specific values, which are highly sensitive to the model type and the influence of technical noise due to imperfect capture of transcripts. From the obtained burst parameter ratios, we deduce how the burst frequency and size vary with cell-cycle phase, thus providing an unprecedented quantitative view of the dynamics of gene expression in mammalian cells.

**Figure 1:**
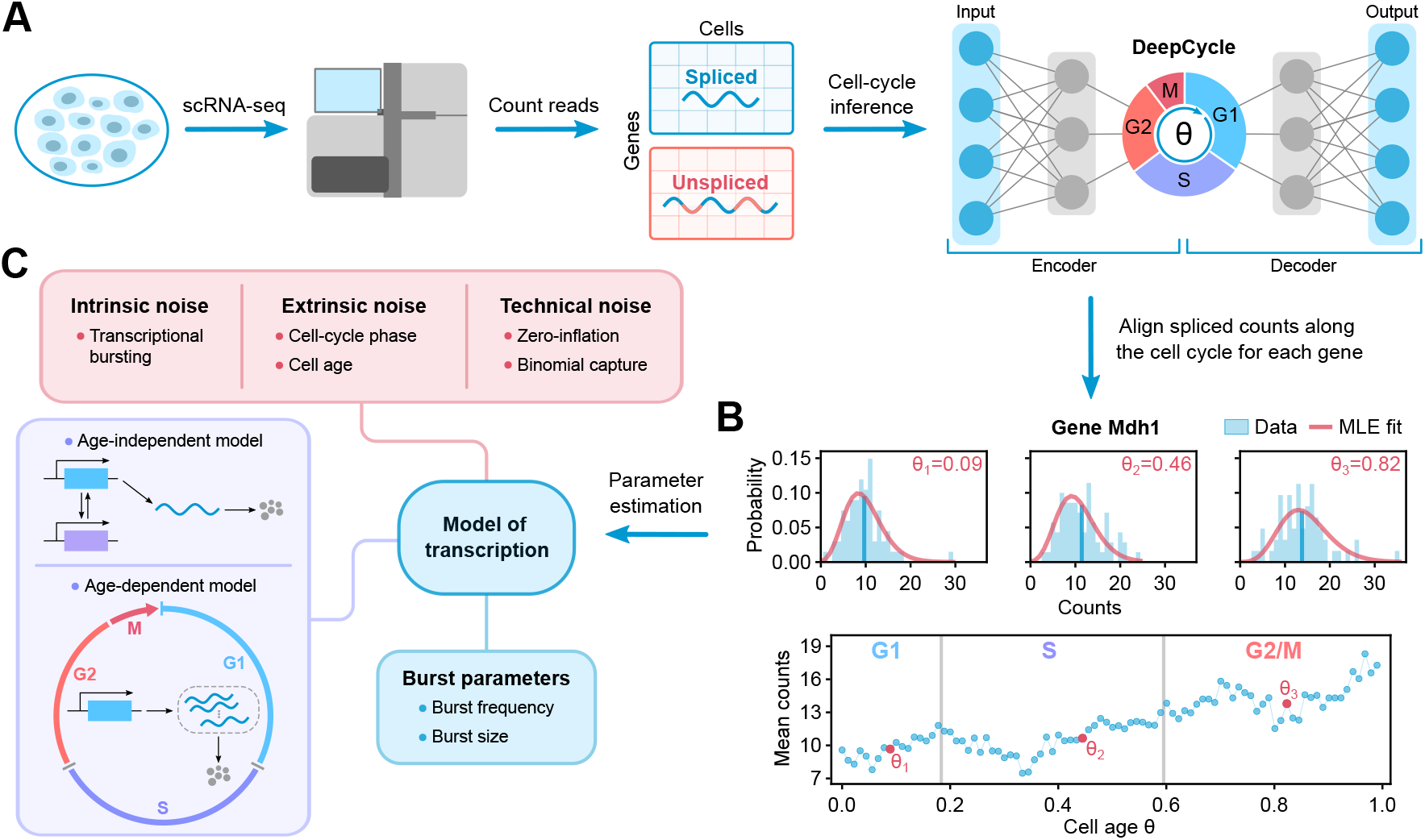
Schematic of the single-cell data acquisition, processing and parameter inference pipeline. **(A)** Our mechanistic analysis is built upon the study of Riba *et al*. [57], which generated a scRNA-seq dataset of over five thousand mouse embryonic stem cells and developed a deep-learning approach called DeepCycle to estimate the cell age and cell-cycle phase for each cell in the population. Specifically, DeepCycle fits the gene expression patterns in the spliced-unspliced space to estimate the cell age *θ* ∈ [0, 1) associated with the relative position of a cell in its progression along the cell cycle from G1 to mitosis. In addition, certain values of *θ* can be linked to transitions between the main cell-cycle phases, and hence all cells in the dataset can be grouped by their cell-cycle phase (G1, S or G2/M). **(B)** An example showing a subset of the processed data for the Prpf8 gene. The bottom graph shows how the mean spliced mRNA counts (average over cells falling in each bin of *θ* values) evolve as the cells go through G1, S, and G2/M phases of the cell cycle. The histograms above show the spliced count distributions for three different values of the cell age *θ*; the vertical blue lines denote the mean values (also indicated by the red circles on the bottom plot). **(C)** In this paper, we use the processed data and the associated cell-cycle information to perform maximum likelihood-based model inference on the experimentally observed (spliced) mRNA count distributions, taking into account intrinsic, extrinsic and technical noise sources. For inference, we use two classes of mechanistic models of stochastic transcription (age-independent models and age-dependent models). As an example, maximum likelihood (MLE) fits for the optimal age-independent model describing the stochastic gene expression of gene Prpf8 are shown by red curves in the upper graphs in (B). The burst parameters (burst size and frequency) are then computed from the optimal model parameters. For technical details see Materials and methods.

## Materials and methods

### Processing of single-cell RNA sequencing data

In our analysis, we use the scRNA-seq dataset of a population of mouse embryonic stem cells with inhibited differentiation generated by Riba *et al*. [57] using the 10x Genomics Chromium platform, containing spliced and unspliced transcript counts for over 5000 cells and more than 10000 genes. Each cell in the population has an associated cell age *θ* (originally referred to as the *transcriptional phase* in [57]) which characterises its position along the cell cycle and can also be used to determine the cell-cycle phase (G1, S or G2/M). The parameter *θ* ∈ [0, 1) is a continuous periodic variable where *θ* = 0 indicates the beginning of the G1 phase of a newly formed cell and *θ* = 1 corresponds to cell division. In the following, we reiterate the main steps of the data processing and downstream analysis published in [57] and discuss how it is reprocessed for the purposes of this paper.

In order to infer the age of each cell, Riba *et al*. developed a neural network-based method called DeepCycle. Their approach is motivated by the concept of RNA velocity, which can be used to estimate the future transcriptional state of a cell from unspliced/spliced mRNA counts [35, 58]. Namely, the authors proposed to order cells along the cell-cycle trajectory by fitting the circular expression patterns of cell cycle-related genes in the unspliced-spliced space using an autoencoder. The data is initially processed using scVelo [36]: the spliced/unspliced counts are normalised to the median of total counts per cell; a nearest-neighbour graph is constructed from principal component analysis on logarithmised counts (using the first 20 principal components); spliced/unspliced counts are smoothed out for each cell by averaging the counts over its 30 nearest neighbours. The next step in the DeepCycle procedure is to identify the cell cycle-related genes that exhibit clearly cycling transcriptional dynamics in the unspliced-spliced space. The z-scored spliced and unspliced counts of the selected genes for each cell comprise the training data for the autoencoder, which learns to denoise and reconstruct the expression patterns of the cycling genes. This way, the single latent variable of the autoencoder, the cell age parameter *θ*, effectively characterises the relative position of a cell along the cell cycle. The variable is then discretised as *θ* ∈ [0, 0.01, …, 0.99] and all cells are assigned to their corresponding bins based on the autoencoder predictions. Finally, specific values of *θ* can be associated with the transitions between the main cell-cycle phases based on the expression of cell-cycle marker genes and the total transcripts per cell — this allows mapping the cells to the G1, S and G2/M cell-cycle phases and to align *θ* so that cell division occurs at *θ* = 1. Specifically, the G1/S and S/G2 transitions occur at *θ* values of 0.26–0.27 and 0.63–0.64 respectively. Although RNA velocity-based methods have various shortcomings [59], Riba *et al*. performed a number of self-consistency checks that establish the validity of DeepCycle for the considered datasets. For instance, the authors verified that the expected expression patterns of marker genes are recovered and also provide complementary validation using flow cytometry analysis.

As the processed mESC dataset published in [57] does not contain the discrete UMI counts needed for our inference and these cannot be easily retrieved, we start from the raw FASTQ data generated by the authors and reprocess it following the same methodology. The reads were aligned to the mouse reference transcriptome (mm10-2020-A) using CellRanger count pipeline (v7.1.0), and the spliced and unspliced transcripts were quantified using velocyto (v0.17.17) [35]. Low quality cells and genes were filtered out using scVelo (v0.2.5) [36]. More specifically, we select genes that are expressed in at least 200 cells and cells that have a total number of UMIs greater than 10000, express more than 3000 unique genes and have a low percentage of mitochondrial genes (under 15%) — the resulting dataset contains 5686 cells and 11544 genes. Next, as described in the paragraph above, we process the data further using scVelo and run DeepCycle to assign the transcriptional phase *θ* ∈ [0, 0.01, … , 0.99] and the cell-cycle phase (G1, S or G2/M) for each cell. Although not exactly equivalent, the obtained phase characterisation is similar both quantitatively and qualitatively to the original processed data in Ref. [57].

In DeepCycle, the transition between mitosis and G1 is defined so that *θ* = 0.99 corresponds to the peak in the average total RNA counts per cell, whereas *θ* = 0 coincides with their immediate drop. Although one may expect to observe a sharp halving in the total counts upon division, this decline is spread over a wider range of *θ* bins assigned to the G1 phase (Figure 3C of [57]) and it remains unclear how many of the cells in this range are misclassified by DeepCycle as belonging to the G1 phase. Therefore, as a precaution, we choose to remove all cells assigned to the [0, 0.09] range of *θ* values and rescale the remaining *θ* ∈ [0.1, 0.11, … , 0.99] into the same range (0–0.99) as *θ* ↦ (*θ* − 0.1) × 0.99*/*0.89. The truncated dataset contains 5294 cells in total, out of which 941 are assigned to the G1 phase, 2240 to the S phase and 2113 to the G2/M phase. Finally, we also remove genes with low expression, i.e., genes that in either G1, S or G2/M phases have a mean transcript abundance below one. The parameter inference is then performed on the spliced mRNA counts of the remaining 3792 genes.

### Mechanistic models of stochastic gene expression

#### Age-independent telegraph, negative binomial and Poisson models

In this paper, we consider the standard model of stochastic gene expression (the telegraph model) [9] and its variants. The telegraph model is defined in terms of the following effective reactions:

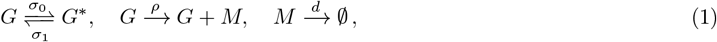

where a gene switches between the active (*G*) and the inactive (*G*^*^) promoter states with rates *σ*_0_ and *σ*_1_, and in the active state produces spliced mRNA (*M* ) with rate *ρ*, which subsequently degrades with rate *d*. Transcriptional bursting is characterised by the burst frequency *f* and burst size *b*, which for the telegraph model are given by *f* = *σ*_1_ and *b* = *ρ/σ*_0_. Note that the burst size is the mean number of mRNA molecules produced when the gene is active.

The chemical master equation (CME) describing the telegraph model can be solved exactly in the steady state for the probability distribution of mRNA counts *m* [1, 9]:

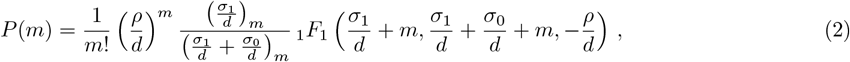

where *(x*)_*m*_ = Γ(*x*+*m*)*/*Γ(*x*) is the Pochhammer symbol and _1_*F*_1_ is Kummer’s (confluent hypergeometric) function. Alternatively, Eq. (2) can be represented by a Beta-Poisson distribution [42, 60]. The rate parameters (*σ*_0_, *σ*_1_, *ρ*) as they appear in the function are divided by the mRNA degradation rate *d*, and hence their values cannot be determined using only static snapshot data without explicit measurements of the degradation rate. Since such measurements are not available, we can only infer the ratios of the rate parameters and the mRNA degradation rate, and hence without loss of generality, in what follows we set *d* = 1.

In the limit *σ*_0_ ≫ *σ*_1_, the gene is mostly inactive and transcription only occurs in short bursts, and the steady-state solution of the telegraph model reduces to a negative binomial distribution:

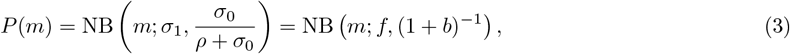

parameterised by the burst frequency *f* = *σ*_1_ and the burst size *b* = *ρ/σ*_0_. Note that this corresponds to the steady-state CME solution of the following reaction network [10]:

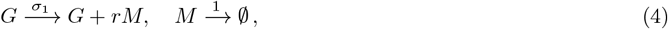

where *r* is a geometrically distributed random variable with mean *b*.

In the limit *σ*_1_ ≫ *σ*_0_, the gene is mostly active and the model reduces to a simple birth-death process:

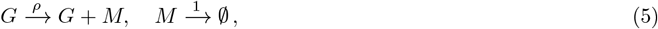

which in steady-state conditions is described by the Poisson distribution *P* (*m*) = Pois (*m*; *ρ*). The burst frequency and burst size cannot be defined in this case as the distribution is characterised by a single parameter, the transcription rate *ρ*, which equals the mean mRNA count.

#### Extending the models to account for zero inflation of scRNA-seq data

Substantial technical noise in scRNA-seq experiments [49, 61] can lead to the presence of excess zero counts in the data. The treatment of such non-biological zeros has been a subject of active debate [50, 62]; although it is commonly taken into account by considering zero-inflated models, a number of recent studies have argued that zero-inflation is largely avoided in UMI count experiments [50–52, 63, 64]. Nevertheless, to allow for more flexibility in our inference procedure, for age-independent models we model zero-inflation explicitly as follows:

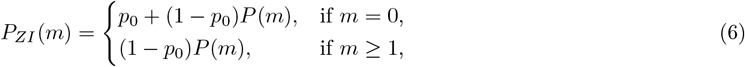

where the parameter *p*_0_ ∈ [0, 1] describes the probability of zero-inflation (fraction of zeros that are non-biological). For age-dependent models, the formula is the same except *P* (*m*) is replaced by *P*_*θ*_(*m*). For parameter inference we specifically consider age-independent telegraph, negative binomial and Poisson models, and their zero-inflated counterparts, i.e. 6 models as illustrated in Figure 1C.

#### Accounting for multiple gene copy numbers

The models considered above describe the transcription from a single allele. However, the mESC dataset that we are analysing in this paper is non-allele-specific, and mouse embryonic stem cells possess two alleles which are replicated during the S phase of the cell cycle. Therefore, we need to consider the combined expression of two gene copies in the G1 phase and four gene copies in the G2/M phase. By assuming that the expression of the alleles is identical and independent from each other, we can model the total count distribution in the G1 phase by convolving the allele-specific count distribution *P* with itself:

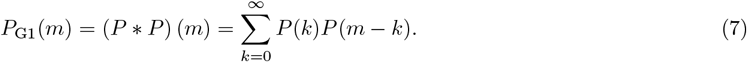

Similarly, the total count distribution in the G2/M phase will be given by the convolution of *P* with itself 4 times:

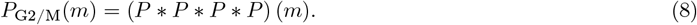

The allele-specific distribution *P* (*m*) is given by the telegraph, negative binomial or Poisson models, or their zero-inflated counterparts, with cell-cycle phase-specific parameters — this is schematically illustrated in Figure 2B. Note that closed-form expressions are known for the convolutions of Poisson and negative binomial distributions, e.g., Pois(*ρ*) * Pois(*ρ*) = Pois(2*ρ*) and NB(*f, p*) * NB(*f, p*) = NB(2*f, p*), which considerably simplifies the numerics.

**Figure 2:**
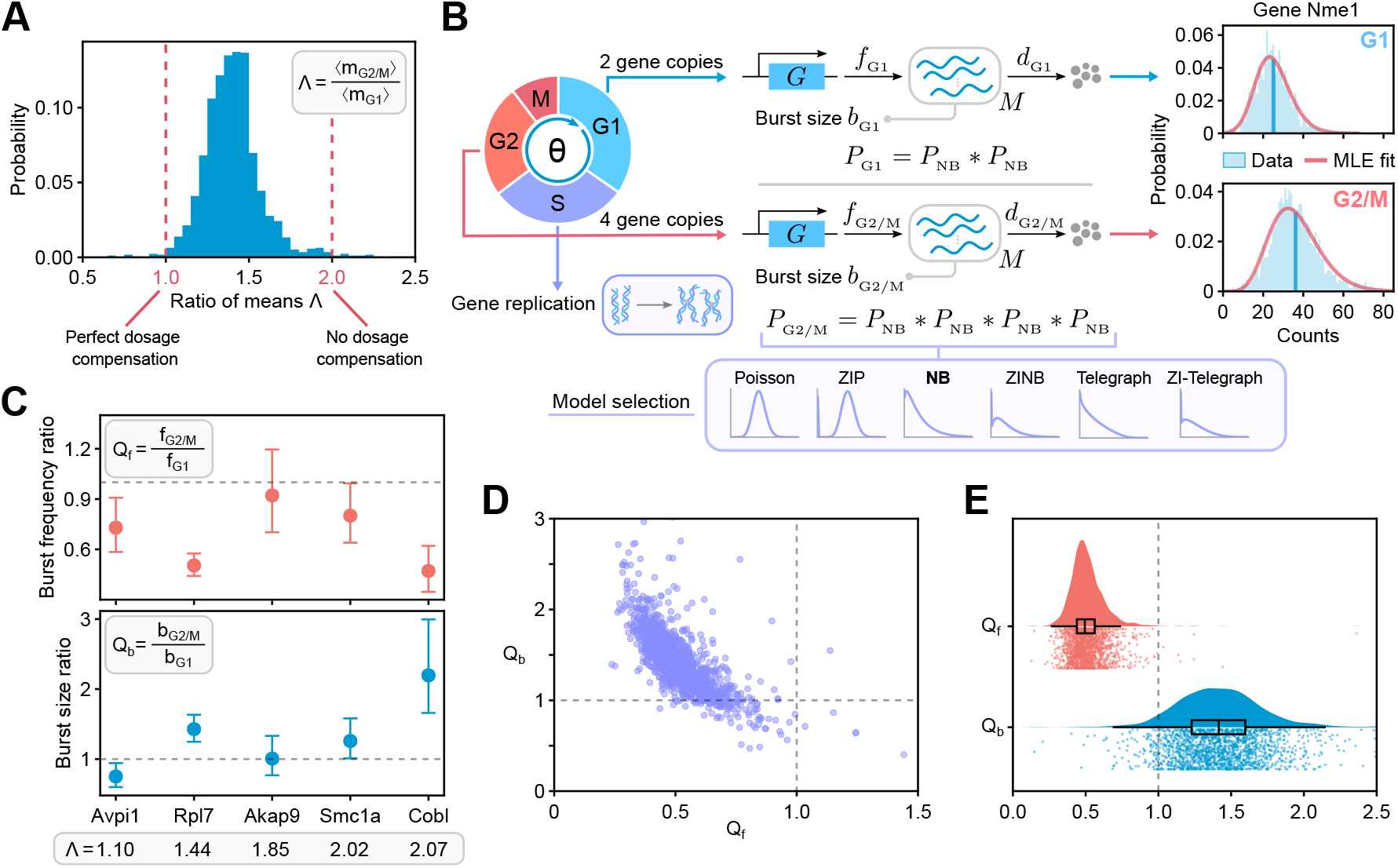
The relationship between cell-cycle phases and the burst parameters. **(A)** Histogram of the ratio of mean mRNA counts in G2/M and G1 phases, Λ = ⟨*m*_*G*2*/M*_⟩ */* ⟨*m*_*G*1_⟩ , for all 1760 bursty genes in the dataset. Ratio of 1 implies a mechanism that perfectly compensates for the doubling of the mRNA number due to the doubling of the gene dosage during replication, while a ratio of 2 indicates the opposite. **(B)** The model fitting and selection algorithm uses six different stochastic models of transcription: Poisson, zero-inflated Poisson (ZIP), negative binomial (NB), zero-inflated negative binomial (ZINB), telegraph and zero-inflated telegraph (ZI-Telegraph) models. One of the 6 models, the one that most successfully describes bursty genes, the negative binomial model, is illustrated in more detail. The rate parameters (normalised by the mRNA degradation rate) are inferred per gene copy. The mRNA distribution in G1, *P*_G1_, is given by a convolution of two negative binomial models *P*_NB_ (with rate parameters having a subscript G1). In the G2/M phase, due to the doubling of gene copy number in the S phase, the mRNA distribution, *P*_G2/M_, becomes a convolution of four negative binomial models (with rate parameters subscripted by G2/M). Plots to the right show the mRNA distributions of gene Nme1 in G1 and G2/M phases and the corresponding best fits using maximum likelihood estimation (MLE fit), where the vertical blue lines indicate the mean values. **(C)** Ratios of the burst frequency per allele (top) and burst size per allele (bottom) in the G2/M and G1 phases, denoted by *Q*_*f*_ and *Q*_*b*_ respectively, for five different genes in the dataset. Error bars indicate the 95% confidence intervals. The grey dashed lines indicate no change in the burst parameter values upon cell-cycle progression. **(D)** Scatter plot of the burst size ratio between G2/M and G1 phases versus the burst frequency ratio for all 1760 genes (each purple dot indicates a single gene). **(E)** Raincloud plots [92] of the burst frequency (red) and burst size (blue) ratios. Each raincloud plot combines a smoothed histogram, a box plot and the jittered raw values of all data points.

#### Extended age-dependent model with bursty transcription

As discussed in the Results section, age-independent (steady-state) gene expression models need to be extended to incorporate the exponential dependence of the mean expression level with cell age *θ*. Here we define the stochastic age-dependent transcriptional bursting model that explicitly incorporates the cell-cycle dynamics (visualised in Fig 3B), and solve it approximately for the mRNA distribution at any cell age *θ*, under the assumption that *θ* is equivalent to the normalised time within a cell cycle of duration *T* , so that *θ* = *t/T* ∈ [0, 1]. The analytic model solution can then be utilised to fit the experimental count data.

**Figure 3:**
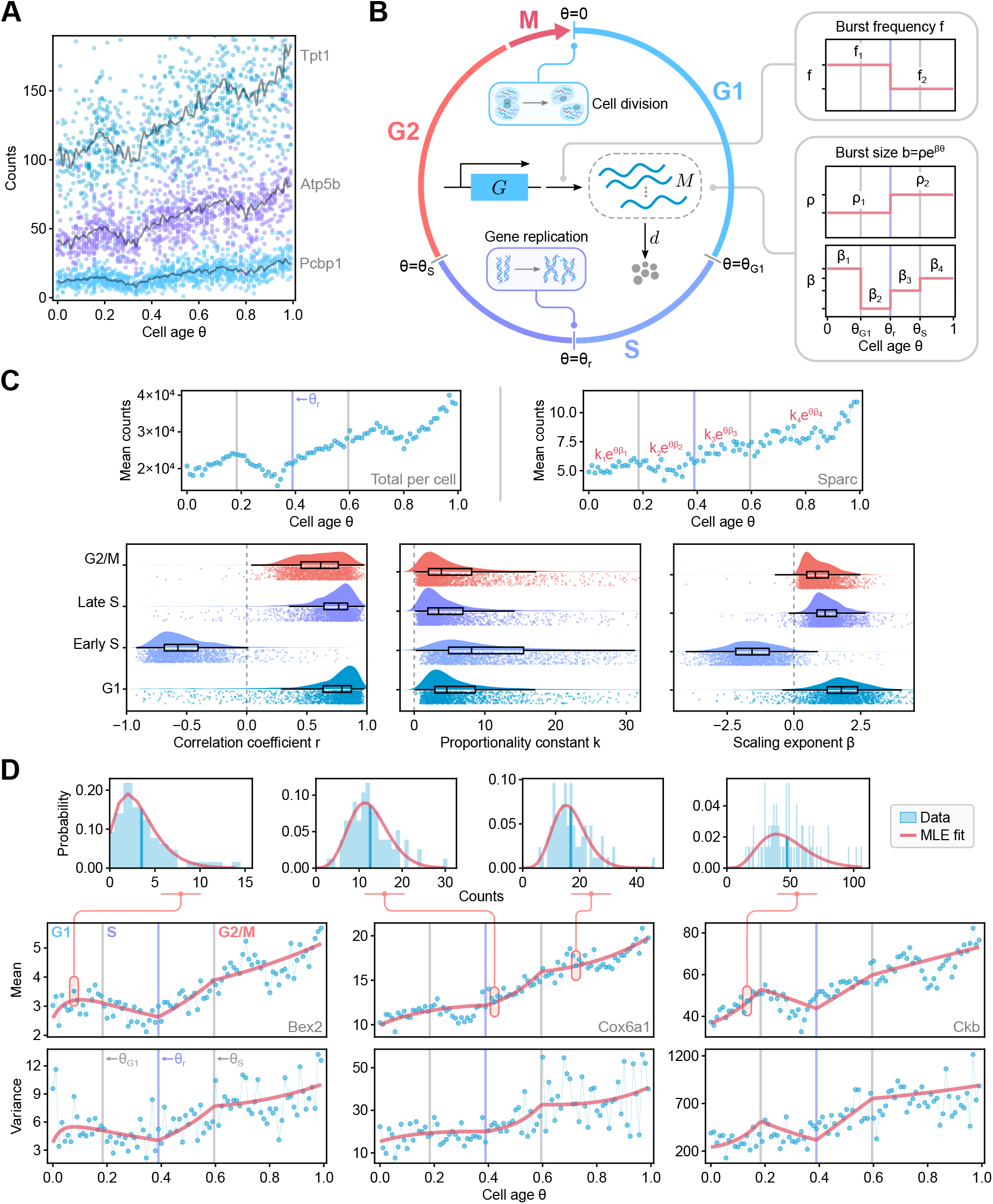
**(A)** Analysis of spliced mRNA counts (from 1000 cells) for 3 genes shows cell-age-dependent expression patterns. Each black line corresponds to the mean mRNA counts (an average over cells falling into each bin of *θ* values). **(B)** Schematic of the age-dependent model describing bursty gene transcription with burst frequency and size that vary with cell age and cycle phase. The model also incorporates the doubling of gene copy number when DNA replication occurs and mRNA dilution when cell division occurs. **(C)** For most genes, there are four main dynamical phases, as shown by a plot of the total mRNA counts versus age (top left). The point half-way through the S phase is denoted by a violet line, while the two grey vertical lines show the G1-S and S-G2/M transitions. An exponential curve *k* exp(*θβ*) is fitted to the data from each gene in each of the 4 phases using non-linear regression (top right). Correlation coefficients and the estimated parameters across phases are visualised for 1760 genes (same as used for Figure 2) using raincloud plots. **(D)** Comparison of the mean, variance and distributions of mRNA counts predicted by the model with parameters inferred using maximum likelihood (red lines; MLE) and the statistics calculated from scRNA-seq data for 3 sample genes.

In the case of transcriptional bursting, the mRNA expression from each allele copy can be described by the following reaction scheme:

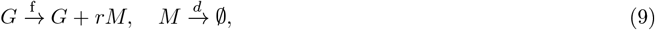

where *r* is an integer sampled from the geometric distribution with mean *b* (the mean burst size), *f* is the burst frequency and *d* is the mRNA degradation rate — this is equivalent to the bursty limit of the telegraph model given by Eq. (4). The age-dependent mean count ⟨*m*⟩(*θ*) is given by the rate equation

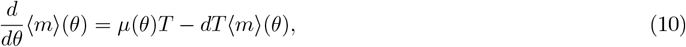

where *µ*(*θ*) is the effective age-dependent mRNA production rate, which equals the product of the burst frequency and the mean burst size at cell age *θ*. In Figure 3C we have demonstrated that the mean mRNA counts in each cell-cycle phase can be well described by an exponential curve *ke*^*βθ*^, pointing out a possible functional form for the effective production rate *µ*(*θ*) in Eq. (10): to enforce ⟨*m*⟩ (*θ*) ∝ *e*^*βθ*^ in our model, it is easy to show that *µ*(*θ*) likewise has to proportional to *e*^*βθ*^, and hence it follows that either the burst frequency or the mean burst size must be proportional to *e*^*βθ*^, with both *β* and the proportionality constant being dependent only on the cell-cycle phase.

To satisfy this parametric constraint, we choose to model the burst frequency *f* as a cell-cycle phase-specific constant and introduce an exponential scaling of the mean burst size with cell age *θ* so that *b* = *ρe*^*βθ*^, where the power *β* is a gene-specific age scaling exponent. Note that the proportionality constant *ρ* in this context has a different interpretation from the transcription rate *ρ* of the standard telegraph model. As discussed in the main text, the choice of functional form here is guided by previous experimental findings. As the burst size *b* is given for each *θ*, we also define the burst size averaged over all cells in a specific cell-cycle phase as:

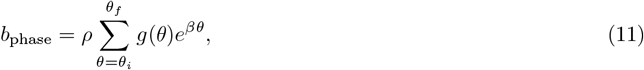

where *θ* varies from *θ*_*i*_ to *θ*_*f*_ and *g*(*θ*) is the distribution of *θ* bins over all cells assigned to that phase (the age distribution in the cell-cycle phase). Note that in the main text we report and discuss the averaged burst sizes in G1 and G2/M, i.e. *b*_G1_ and *b*_G2/M_, where the parameters *ρ* and *β* in Eq. (11) are the piecewise constants *ρ*_1_ and *β*_1_ for cells in G1, and *ρ*_2_ and *β*_4_ for cells in G2/M.

We construct the age-dependent model in a piecewise manner in order to account for the extrinsic noise due to the cell-cycle phase and the doubling of gene copy number upon DNA replication at *θ* = *θ*_r_ (see Fig 3B for a visual summary). Namely, we assume that the allele-specific burst parameters *f* and *ρ* are given by the respective constants *f*_1_ and *ρ*_1_ before replication, and *f*_2_ and *ρ*_2_ after replication. Furthermore, the parameter *β* that characterises the exponential scaling of transcription with *θ* is piecewise parametrised by four constants: *β*_1_ in the G1 phase for *θ* = [0, *θ*_G1_], *β*_2_ in the S phase before replication for *θ* = [*θ*_G1_, *θ*_r_], *β*_3_ in the S phase after replication for *θ* = [*θ*_r_, *θ*_S_], and *β*_4_ in the G2/M phase for *θ* = [*θ*_S_, 1], where *θ*_G1_ and *θ*_S_ are the transition time points between the G1/S and S/G2 phases respectively (as determined by DeepCycle). In summary, the age-dependent production rate in Eq. (10) is equal to

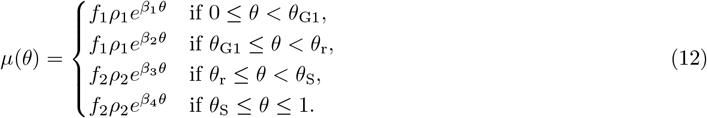

Note that the parameter *β* is segmented into four parts in order to introduce more model flexibility and better fit the non-monotonic trend of the mean counts with respect to cell age *θ* observed in the S phase. While we could similarly consider the parameter *ρ* to be a different constant in each of the four phases, we only allow it to take two values (one prior to and another post-replication). This choice is motivated by the fact that *β* is the parameter which most strongly varies between the phases (bottom row of Figure 3C), and by the practical desire to reduce the number of model parameters that need to be estimated for each gene.

To inform the model we use a fixed cell cycle duration *T* = 13.25 h, experimentally determined as the median cell cycle length of mESCs in the pluripotent ground state [65] (grown in 2i + LIF medium, similarly to the data generated by Riba *et al*. [57]). Gene-specific degradation rates *d* are obtained from an mRNA decay database for differentiating mESCs [66], discussed in more detail in the Gene filtering section. Although gene-specific DNA replication times *θ*_r_ could in principle be extracted from the existing DNA replication timing profiles for mESCs [67], to our knowledge these are not readily available in experimental literature without requiring additional processing and technical considerations. Hence, for simplicity, we fix the replication time for all genes in the very middle of the S phase at *θ*_r_ = (*θ*_G1_ + *θ*_S_) */*2. In Supplementary Text S3, we argue in favour of this assumption by performing parameter inference using an alternative age-dependent model formulation that avoids setting a fixed replication time but still leads to qualitatively similar observations on the transcriptome level.

Using Eqs. (10) and (12), we can derive piecewise differential equations that describe the time evolution of the mean mRNA counts due to all (independent) alleles in each of the four cell-cycle phases:

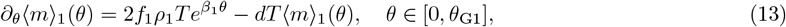

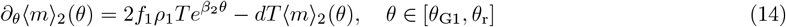

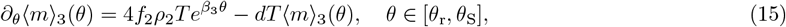

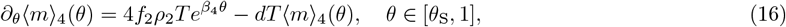

where the additional factors of 2 and 4 stem from the number of alleles in each cell-cycle phase and the subscript *i* in ⟨*m*⟩_*i*_(*θ*) denotes the mean mRNA in the *i*-th cell-cycle phase. There are four boundary conditions:

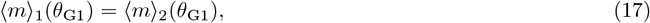

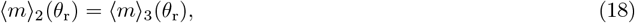

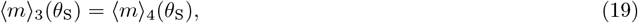

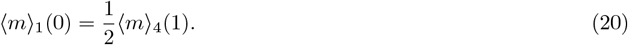

The first three boundary conditions ensure that the mean mRNA count is continuous as the cell-cycle phase progresses from G1 to G2, while the last boundary condition models dilution due to the binomial partitioning of mRNA molecules at cell division with probability 1*/*2, i.e. on average only half of the mRNA molecules from generation *i* are present when the next generation *i* + 1 starts. Note that here we consider the steady-state growth conditions, i.e. enough time has passed such that the probability that a cell of age *θ* has a given number of mRNA molecules is independent of which generation it belongs to [68] — this means that there is no generation index in our equations.

Similarly from the chemical master equation for reaction scheme (9) we can derive equations for the variance of the mRNA counts due to all (independent) alleles in each cell-cycle phase:

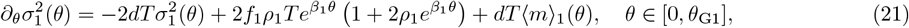

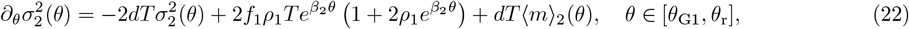

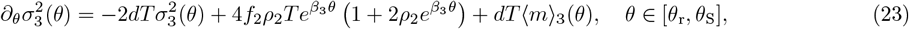

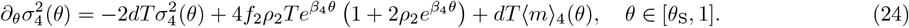

The boundary conditions satisfied by these ordinary differential equations (ODEs) are:

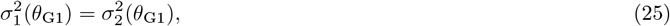

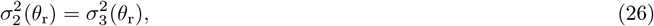

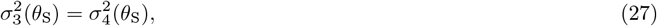

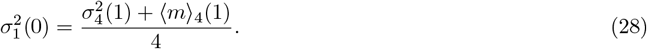

As for the mean, the first three conditions stem from continuity of the variance at the transition points between the cell-cycle phases and the gene replication point, while the last boundary condition is due to binomial partitioning of the mRNA molecules at division with probability 1*/*2. This last condition can be derived from the equation linking the mRNA distribution at the beginning and the end of the cell cycle [68]:

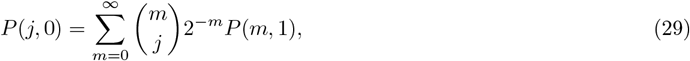

where *P* (*i, θ*) is the probability of observing *i* mRNA molecules at time *θ*.

We can solve the ODEs for the means (given by Eqs. (13)-(16)) and variances (Eqs. (21)-(24)) with the corresponding boundary conditions (Eqs. (17)-(20) and (25)-(28)) using Mathematica. As the obtained expressions are lengthy, we omit them from the manuscript — the complete Mathematica notebook is given in the associated GitHub repository (see Data Availability Statement).

Finally, we assume that the distribution of mRNA counts at any cell age *θ* is well approximated by a negative binomial distribution whose first and second moments agree with the exact moment solutions derived above:

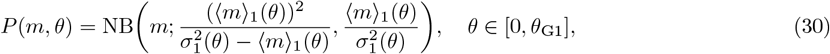

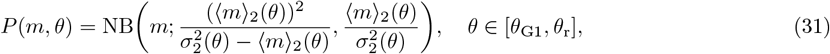

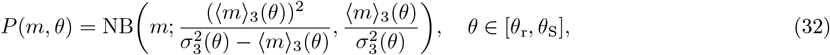

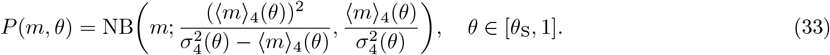

Note that this approximation is based on previous work which showed that in complex models of bursty expression including cell division, DNA replication and other phenomena, the time-dependent distributions were generally well approximated by the negative binomial distribution [69]. This assumption is further supported by the model selection results using steady-state age-independent models, revealing that bursty genes are in almost all cases optimally fit by the negative binomial model. Moreover, in Supplementary Text S2 (and Supplementary Figure S2) we benchmark the performance of our negative binomial approximation using synthetically generated count data, further validating its accuracy.

### Parameter inference

We use Julia [70] for parameter inference and subsequent analysis, utilising Distributions.jl [71] in the implementation and Makie.jl [72] for the visualisation of results, together with a number of numerical optimisation packages listed below.

#### Maximum likelihood estimation

Let ***m*** = (*m*_1_, … , *m*_*n*_) denote a vector of spliced mRNA counts for one gene over *n* cells in a specific cell-cycle phase. The likelihood of observing the data ***m*** for a model with a count distribution *P* ( ·|***ξ***) given a set of parameters ***ξ*** is defined by

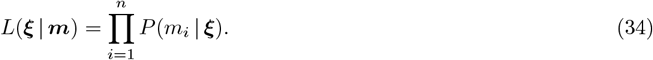

In practice, to find the maximum likelihood estimate (MLE) of the model parameters, ***ξ***^*^, we minimise the negative log-likelihood:

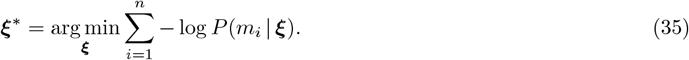

Note that for the age-dependent model we have to maximise the likelihood by explicitly taking into account the age *θ* of each cell. Over the entire cell cycle, with the associated *θ* bins in the range from *θ*_*i*_ = 0 to *θ*_*f*_ = 0.99, the optimal parameters will be given by:

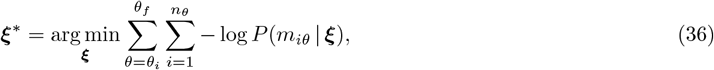

where *n*_*θ*_ is the number of cells in a given *θ* bin and *m*_*iθ*_ is the mRNA count in cell *i* with the same *θ* value.

We fit the steady-state age-independent gene expression models to the scRNA-seq data for all 3792 genes using maximum likelihood. Note that each model here refers to the total count distribution given by a *k*-fold convolution of the allele-specific count distribution with itself, where *k* is the number of gene copies in a given cell-cycle phase — throughout the paper we report the burst parameters per gene copy. We compute the count distributions explicitly (as given in the section on Mechanistic models of stochastic gene expression) to obtain the likelihood of each model. Although some studies have highlighted the computational challenges of evaluating the steady-state solution of the telegraph model involving a confluent hypergeometric function [42], we found our direct implementation using log-probabilities to be sufficiently numerically stable and more efficient than the corresponding Beta-Poisson formulation using Gaussian quadratures [43, 73]. Note that parameter estimation using the age-dependent gene expression model is performed only for 1351 bursty genes, as covered in the Gene filtering section.

To minimise the negative log-likelihood, we initially perform a global optimisation using the adaptive differential evolution optimiser with radius limited sampling, as implemented in BlackBoxOptim.jl [74]. To ensure convergence, we then use the candidate solution as the initial value for a local search algorithm — here we employ the BFGS optimiser implemented in Optim.jl [75]. The global and local optimisation steps are both terminated after 60 seconds, albeit they generally converge much faster for the majority of genes. Although the optimiser is quite sensitive to the initial condition and may get stuck in a bad local minimum and fail to converge in the designated times for the telegraph-like steady-state models or the age-dependent model due to parameter unidentifiability, we found that simply restarting the inference procedure for the problematic genes typically resolved the issue. For this reason, we automate the procedure further by rerunning the inference for all genes with a different random initial condition for the global optimiser 5 times using the age-independent models (10 times for the age-dependent model) and choosing the fit with the best likelihood value.

The optimisation is performed in linear, log- or logit-transformed parameter space with model-specific parameter bounds. Note that for the Poisson model the solution is simply given by the sample mean, which in the zero-inflated case is constrained as *ρ* ∈ [ −9, 7] on the log-scale. The search space for the negative binomial model, NB(*f, p* = (1 + *b*)^−1^), is constrained to *f* ∈ [ −10, 10] on the log-scale and *p* ∈ [ −30, 30] on the logit-scale (as *p* is a probability from zero to one). For the telegraph model, we fix *σ*_0_, *σ*_1_, *ρ* [ 9, 7] on the log-scale. The zero-inflated counterparts of the mentioned models have exactly the same parameter constraints, and we set the zero-inflation probability to *p*_0_ ∈ [ −30, 30] on the logit-scale. The search space for the piecewise parameters characterising the age-dependent model is constrained to *f*_1_, *f*_2_, *ρ*_1_, *ρ*_2_ ∈ [ −7, 7] on the log-scale and *β*_1_, *β*_2_, *β*_3_, *β*_4_ ∈ [−100, 100] on the linear scale, in order to allow for the negative exponential scaling of transcription with the cell age *θ* observed in the S phase.

#### Steady-state model selection

We determine the optimal steady-state model for each gene using the Bayesian information criterion (BIC):

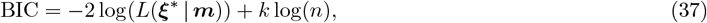

where *k* is the number of model parameters, *n* is the number of data points (cells) and *L*(***ξ***^*^| ***m***) denotes the maximised likelihood function for the data vector ***m***. Given two models *M*_1_ and *M*_2_, the difference between their respective BIC values BIC_1_ and BIC_2_, Δ = BIC_1_ − BIC_2_, reflects the strength of evidence in favour of *M*_2_ over *M*_1_, with Δ *>* 10 constituting “very strong” evidence for model *M*_2_ over *M*_1_ [76, 77]. We utilise this criterion to iteratively compare six different models (from the simplest to the most complex): Poisson, zero-inflated Poisson (ZIP), negative binomial, zero-inflated negative binomial, telegraph and zero-inflated telegraph (ZI-telegraph) models.

We start by computing the difference in the BIC values between the Poisson model and all the other candidates, i.e. Δ_1_ = BIC_Poisson_ − BIC_ZIP_, … , Δ_5_ = BIC_Poisson_ − BIC_ZI-Telegraph_. If all five Δ values are smaller than 10, we pick the Poisson model as the optimal choice; otherwise, we proceed to compute the difference in the BIC values between the zero-inflated Poisson and the remaining (more complex) models, repeating the procedure until the best candidate is found. This allows us to filter out the models with practically unidentifiable parameters and select the simplest gene expression model that still provides a good fit to the mRNA count data. We verify the robustness of our model selection procedure in Supplementary Text S1 (also see Supplementary Figure S1).

The model selection is performed for all 3792 genes in the dataset in both G1 and G2/M phases. As the burst frequency and size parameters cannot be extracted from the Poisson and zero-inflated Poisson models, all genes that either in G1 or in G2/M are best fit by the Poisson or zero-inflated Poisson models are excluded from further analysis. After this filtering step applied to the age-independent models, we are left with 1760 genes that in both cell-cycle phases are best fit by either the negative binomial, zero-inflated negative binomial or telegraph models. Namely, we find that the overwhelming majority of the remaining genes are best fit by the negative binomial model; the zero-inflated negative binomial is the optimal model for 22 genes in G1 and 15 genes in G2/M; the telegraph model fits only 1 gene in G1 and 2 genes in G2/M; and the zero-inflated telegraph model was not found to be optimal for any genes.

#### Gene filtering for the age-dependent model

Following the model selection performed for (steady-state) age-independent models, we are left with 1760 bursty genes that are mostly best fit by the negative binomial model. To proceed with the analysis of these bursty genes using the age-dependent model, we perform a series of gene filtering steps prior to and following parameter inference, as outlined below.

The age-dependent model requires us to provide an explicit mRNA degradation rate for each individual gene. We use the mRNA decay rates from Supplementary Table S2 [66] obtained for mESC lines MC1 (129S6/SvEvTac) and MC2-B6 (C57BL/6J) grown in the presence of Leukemia Inhibitory Factor (LIF): if the gene-specific decay rate is negative for MC1 due to experimental constraints, we use the decay rate obtained for MC2 cells if it was found to be positive (as the observed correlation between the strains on average is high), and consider the decay rate to be unknown otherwise. Out of the 1760 initial bursty genes we thus discard 334 genes that do not have an associated mRNA degradation rate.

Next, we filter out 75 genes that display a negative correlation between the mean mRNA transcription and cell age *θ* either in the G1 or G2/M phases. Most genes in this category tend to switch on in the S phase or display a downward trend in the mean counts in the G1 phase with respect to *θ* — these genes are outliers in terms of their phase-specific transcriptional profiles, and hence for simplicity we choose to discard them.

After the two filtering steps, we fit the age-dependent model to mRNA count data for the remaining 1351 bursty genes using maximum likelihood. We then filter out 291 genes with poor model fits, where the predicted mean expression in the G1 or G2/M phases negatively correlated with cell age *θ*, whereas the observed count data showed the opposite trend.

Lastly, we compute the coefficient of determination *R*^2^ for the predicted mean and variance of age-dependent model fits over the entire cell cycle to identify any other extreme outliers. We define the coefficient of determination as 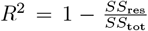, where *SS*_res_ and *SS*_tot_ stand for the residual sum of squares and the total sum of squares, respectively. We choose to discard 16 genes that have negative associated *R*^2^ values, which can occur when fitting a non-linear model without an intercept term and which usually indicate bad model fits [78].

Following all filtering steps we are left with 1044 genes, which we investigate further in the main text using the age-dependent model.

#### Confidence intervals

To quantify the uncertainty in the best fit model estimates, we compute the 95% confidence intervals for each model parameter using profile likelihood [79, 80]. If we partition a vector of model parameters ***ξ*** into a parameter of interest *ϕ* and a vector of nuisance parameters ***ψ***, i.e. ***ξ*** = (*ϕ*, ***ψ***), the profile likelihood of the parameter *ϕ* given a set of observations ***m*** can be defined as:

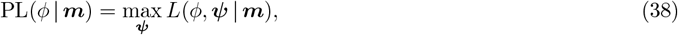

where we maximise the likelihood keeping *ϕ* fixed. It can be shown that the approximate 100(1 − *α*)% confidence interval for *ϕ* is given by [79]:

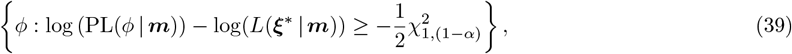

where *L*(***ξ***^*^ |***m***) is the maximised likelihood (with all parameters allowed to vary) and 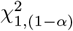 is the (1−*α*) quantile of the *χ*^2^ distribution with one degree of freedom.

We can generalise the approach above to compute the confidence intervals for some function of the model parameters, *g*(***ξ***), by defining the prediction profile likelihood [80, 81]:

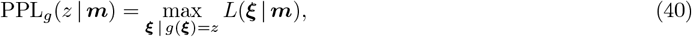

where we maximise the likelihood under the constraint that *g*(***ξ***) equals the value *z*. Similarly to the confidence interval for a single model parameter, the 100(1 − *α*)% confidence interval for the set of predictions *g*(***ξ***) = *z* is given by:

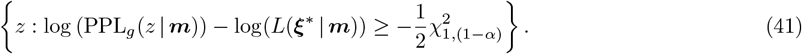

We can also find the confidence intervals for functions of the parameters of multiple different models by considering their joint likelihood. For example, consider the ratio of the burst frequency between the G2/M and G1 phases, *Q*_*f*_ = *f*_G2/M_*/f*_G1_ (examined in Figure 2C). Let ***m***_G1_ and ***m***_G2/M_ be the vectors of mRNA counts in G1 and G2/M respectively, and similarly define ***ξ***_G1_ and ***ξ***_G2/M_ as the phase-specific model parameters. The joint likelihood is then given by

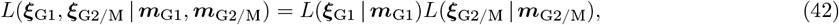

and the prediction profile likelihood can be written as

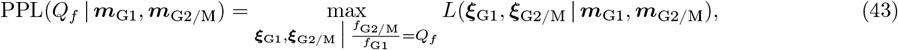

so that the confidence interval for *Q*_*f*_ is similar in form to Eq. (41).

The outlined approach is used throughout the paper to compute the confidence intervals for any considered quantity *z*. We implement this numerically by simply increasing (decreasing) *z* in small steps starting from its maximum likelihood value, and recomputing the prediction profile likelihood at each step until the threshold defining its upper (lower) confidence interval limit is reached. We perform the constrained optimisation using the Ipopt algorithm [82] through the MetaOptInterface.jl [83] and Optimization.jl packages. The parameter bounds for each model are the same as previously used for maximum likelihood estimation.

### Binomial downsampling of count data

In this section, we describe the implementation details of our binomial downsampling strategy for scRNA-seq data, which is analysed in the main text. In brief, we simulate the true mRNA count data and downsample it based on a binomial mRNA capture model [84, 85] (visualised in Figure 4D left) assuming that there is a finite capture probability *p* of detecting each mRNA molecule in each cell in an scRNA-seq experiment (where *p* varies from cell to cell according to a probability distribution).

**Figure 4:**
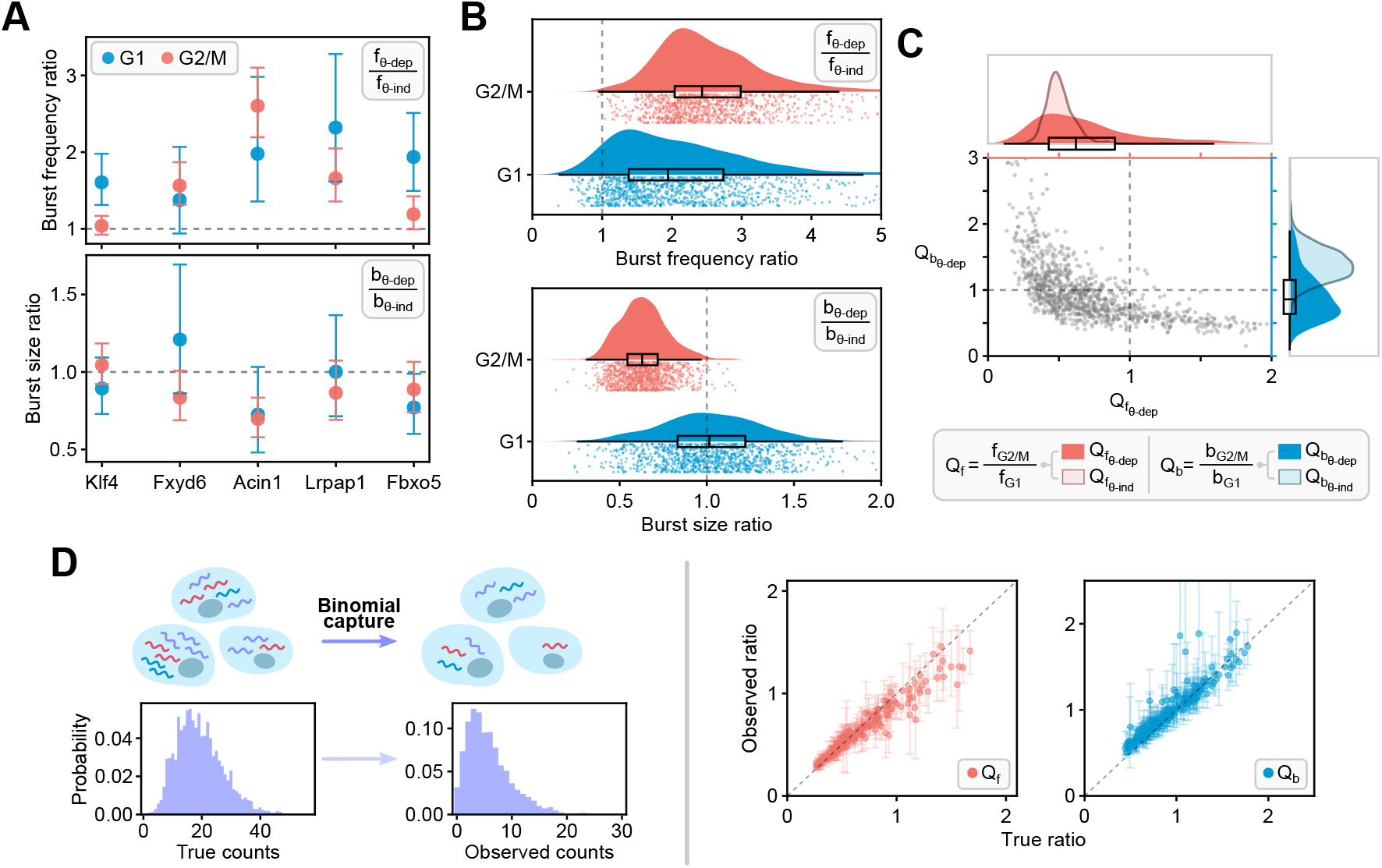
Effect of extrinsic noise due to age-dependent transcription rates and technical noise on the burst parameter inference. **(A)** Burst frequency (top) and burst size (bottom) ratios between *θ*-dependent (*θ*-dep) and *θ*-independent (*θ*-ind) model estimates for five different genes in the dataset. The ratios are inferred both for G1 (blue) and G2/M (red) phases. The error bars show 95% confidence intervals. The grey dashed lines indicate no change in the parameter values between the two cell-cycle phases. **(B)** Raincloud plots of burst frequency (top) and burst size (bottom) ratios between *θ*-dependent and *θ*-independent model estimates for all genes, inferred for G1 (blue) and G2/M (red) phases separately. **(C)** Scatter plot of the burst size ratio, 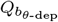, versus the burst frequency ratio, 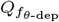, between the G2/M and G1 phases for all 1044 genes (each grey dot indicates a single gene) estimated with the *θ*-dependent bursty model. The smoothed histogram on the right (in blue) represents the distribution of burst size ratios, and the distribution of burst frequency ratios is given on the top (in red). The equivalent distributions of burst frequency and burst size ratios between G2/M and G1 obtained using the *θ*-independent model (shown in Fig 2D), i.e. 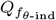 and 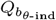, are superimposed as transparent smoother histograms with darker contours for comparison. **(D)** Left: Illustration of the binomial model of mRNA capture. In a sequencing experiment, only a fraction of the original mRNA counts will be detected due to the finite capture probability *p* that varies between cells. Right: scatter plot of the burst frequency (red) and burst size (blue) ratios between the G2/M and G1 phases estimated by fitting the age-dependent model to the true count data and the corresponding downsampled (observed) data, obtained by simulating binomial transcript capture (see text for details).

The mESC library published by Riba *et al*. [57] is generated using the 10x Genomics Single Cell 3’ Reagent Kit (v3), which reportedly (according to the manufacturer) captures 30-32% of transcripts per cell, and hence we assume that the mean capture efficiency is given by ⟨*p*⟩ = 0.3. To address the heterogeneity in capture efficiencies amongst cells, we add small noise by assigning the capture efficiency *p* for each cell to be a random number drawn from Beta(30, 70) distribution with mean ⟨*p*⟩ and coefficient of variation CV_*p*_ ≈ 0.15. Although the assumed noise model is quite simplistic, more accurate assignment of capture efficiencies would require spike-in data (unavailable for this mESC experiment) and careful statistical analysis to correct for other potential cell- or gene-specific biases — to our knowledge, studies focusing on mRNA capture efficiency specifically are quite limited and make additional modelling assumptions [85, 86], while more relevant analyses for the 10x Genomics v3 Kit do not appear to be publicly available.

We consider 200 genes randomly chosen from the subset of all bursty genes fit by the age-dependent model whose burst frequency and burst size ratios between the G2/M and G1 phases, i.e. *Q*_*f*_ and *Q*_*b*_ values, lie in their respective 5-95 percentile ranges (Figure 4C). For each gene, we use the best-fit age-dependent model and rescale its parameters *ρ*_1_ and *ρ*_2_ by the inverse of the average capture efficiency ⟨*p*⟩, thus rescaling the mean burst size so that downsampled counts resemble the experimental data. The observed mRNA count *x* for each gene in each cell at a certain cell age *θ* is obtained by sampling the true count *y* from the age-dependent negative binomial mRNA distribution of the rescaled model and subsequently downsampling *y* so that *x* ∼ Binomial(*y, p*). By repeating this process for each of 5294 cells, we construct a downsampled dataset equivalent in structure to the original data with the same number of data points in each *θ* bin. We use this procedure to generate 100 of such datasets and fit them to the age-dependent model. In Figure 4D right, we plot the observed median burst parameter ratios between the G2/M and G1 phases with their respective interquartile ranges for the 200 selected genes against the ground-truth estimates fit to the original count data (for further discussion see Results).

## Results

### Inference of cell-cycle dependent burst parameters using age-independent models

To understand whether the burst parameters vary across the cell cycle, we calculate the mean mRNA counts of each gene using data from all cells that are in G1 (⟨*m*_G1_⟩) and similarly using data from all cells in G2/M (⟨*m*_G2*/*M_⟩). In Figure 2A, we show the distribution of the ratio of the means (Λ = ⟨*m*_G2*/*M_⟩ */* ⟨*m*_G1_⟩ ). Note that the mass of the distribution is concentrated between Λ = 1 and Λ = 2 with a median Λ = 1.40. The case Λ = 2 is expected if there is no change in the rate parameters from G1 to G2/M, since the doubling of the gene copies would simply imply a corresponding doubling of the mean mRNA count. In contrast, the case Λ = 1 is what can be described as perfect gene dosage compensation, in the sense that the rate parameters have changed in such a way that the mean mRNA count remains unaltered as the cell cycle progresses. For the vast majority of genes, Λ is neither 1 nor 2, implying that there is a change in the rate parameters between the two cell-cycle phases but it is insufficient to perfectly compensate for the doubling of the gene copy number during DNA replication. Hence we conclude that partial dosage compensation is the norm across the transcriptome.

Given the foregoing conclusion that for most genes there is a change in the rate parameters as the cell cycle progresses, we next infer the burst parameter values in each phase. We assume that in each of the two phases, the distribution of spliced mRNA counts is well described by the steady-state distribution of a stochastic model of gene expression describing transcription from *N* independent promoters, where *N* = 2 in the G1 phase and *N* = 4 in the G2/M phase. The mRNA count distribution of this model is obtained by convolving the steady-state mRNA distribution of a stochastic model of gene expression for one promoter with itself *N* times; this is equivalent to assuming that expression from individual gene copies is independent of each other, a property of many eukaryotic genes [22, 34, 43] (this assumption can be avoided if the scRNA-seq data is allele-specific [43]). We refer to this type of model as an age-independent model to emphasise that here we assume that all cells within each cell-cycle phase, independent of their cell age, have the same burst size and frequency; the burst parameters only change with cell-cycle phase. Using the model count distribution and the transcript count data for each cell, we use the method of maximum likelihood to estimate the rate parameters (normalised by the mRNA degradation rate) in each cell-cycle phase. We fit six different gene expression models (telegraph model, negative binomial model, Poisson model and their equivalents with zero-inflation) separately to G1 and G2/M count data from 3792 genes that satisfied quality control criteria, and then select the optimal model using the Bayesian Information Criterion. More details regarding the age-independent models, parameter inference and model selection are provided in Materials and methods.

We note that in our repertoire of models we have not included models more complex than the telegraph model. Some (non-sequencing) studies that can directly measure the distribution of the times spent in the active and inactive gene state suggest the three-state model as the optimal model of gene expression in mammalian cells [2, 87]. Models that are more complex than the (two-state) telegraph model can be fit to snapshot scRNA-seq data to estimate the burst parameters [88]. However, as the steady-state mRNA count distributions of the three-state and the telegraph models can often equally well fit snapshot count data from a few thousand cells [89–91], a model selection algorithm is unlikely to select a three-state model over a two-state model because the former has more parameters than the latter; this issue is particularly pronounced for bursty gene expression, since both models reduce to the negative binomial model in this case.

We find that 55% genes are best fit by a Poisson or a zero-inflated Poisson model which describes non-regulated expression from an active gene state. The rest of the genes are best fit by models that implicitly or explicitly assume gene state switching between active and inactive states, and hence we call them bursty genes since their expression occurs in bursts when the gene is active. Most of these genes are fit by the negative binomial model (44%) and very few (1%) by the rest of the models. The model that most successfully describes bursty genes, the negative binomial model (also called the bursty model), is illustrated in Figure 2B. Note that in these age-independent models, the burst parameters are normalised by the mRNA degradation rate (Materials and methods).

For each of the bursty genes, of which there are 1760, we calculate *Q*_*f*_ = *f*_G2*/*M_*/f*_G1_ and *Q*_*b*_ = *b*_G2*/*M_*/b*_G1_. These are the ratios of the burst frequency and burst size (for each gene copy) in the two phases, respectively. In Figure 2C we show these ratios for 5 genes. Note that there appears to be a tendency for the burst frequency in the G1 phase to be greater than that in the G2/M phase, and the opposite tendency is observed for the burst size. In Figure 2D we show the distributions of *Q*_*f*_ and *Q*_*b*_ for all genes, confirming that for most genes there is an attenuation of the burst frequency and a corresponding amplification of the burst size as the cell cycle progresses. Specifically, the distribution of *Q*_*f*_ has a median of 0.49 with first and third quartiles of 0.44 and 0.56, while the distribution of *Q*_*b*_ has a median of 1.41 with first and third quartiles of 1.23 and 1.60.

Note that as a first step, we have fitted age-independent models to cell-cycle phase-specific data because their steady-state mRNA count distributions are all known in closed form and they are straightforward to fit using the method of maximum likelihood. However, these models suffer from two major disadvantages: (i) they implicitly assume that the cells in each of the cell-cycle phases are identical, i.e. the burst parameters within each cell-cycle phase do not vary from cell to cell, and hence cannot be age-dependent; (ii) they assume that shortly after a cell transitions from one cell-cycle phase into another, a new steady state is quickly reached. Regarding (i), as mentioned in the Introduction, there is evidence from smFISH studies of extrinsic noise within the cell cycle for various types of mammalian cells; however, we do not know of studies specific to mESCs. Regarding (ii), rough estimates can be made to see whether the steady-state assumption generally holds. The median mRNA half-life is estimated to be 7.1 hours [66], while the median cell-cycle duration is estimated to be 13.25 hours for mESCs with blocked differentiation [65]. Since the median half-life is comparable to or longer than the time spent in each cell-cycle phase, for many mRNA species it is unlikely that a steady state is reached within each cell-cycle phase. Evidence supporting this hypothesis is shown in Figure 3A and C (top left plot): the number of spliced mRNA counts tends to increase with cell age *θ* within the cell cycle. It is hence possible that due to the aforementioned limitations of age-dependent models the inference procedure has introduced biases in parameter estimation.

### Constructing a sophisticated cell-age and cell-cycle phase-dependent model

A schematic of a cell-age dependent model of bursty gene expression that overcomes the limitations of previous models is shown in Figure 3B. At its core, the model assumes that the transcription of mRNA *M* from each allele copy *G* occurs in bursts at an associated burst frequency *f*, where the number of molecules produced in a burst is a geometrically distributed variable with the mean burst size *b*, and the mRNA decays at a constant rate *d* [14]. We assume that all parts of the cell cycle are proportional to the total division time, i.e. the stretched cell-cycle model proposed in [93, 94]. Specifically, by letting *θ* = *t/T* where *T* is the median cell-cycle duration time, i.e. assuming that the cell age is equivalent to the normalised time within the cell cycle, it follows that the cell cycle starts at *θ* = 0, changes from G1 to S phase at *θ* = *θ*_G1_, from pre-DNA replication to post-DNA replication at *θ* = *θ*_r_ and from S to G2/M phase at *θ* = *θ*_S_. Note that since we do not know the precise cell age at which each gene replicates, for simplicity, for all genes we choose *θ*_r_ to lie exactly in the middle of the S phase (while this assumption may appear rough, as we show later on, it has only a minor influence on the inference results). Furthermore, it is assumed that the number of gene copies changes from 2 to 4 upon DNA replication, and the number of mRNA copies are randomly partitioned with probability 1*/*2 between the two daughter cells upon (symmetric) cell division. What remains for full model specification is to devise functions that describe how the burst size and frequency change as a function of *θ* within the four ranges of the cell-cycle phase in our model (G1, S phase prior to replication, S phase post-replication and G2/M).

To obtain some insight into how we can choose these functions in a manner that is biologically realistic, we plot the total mRNA counts versus age (top left corner of Figure 3C) — the plot shows the main four dynamical phases (increasing/decreasing/increasing/increasing) which correspond to G1, early S, late S and G2/M phases. It is unclear what causes the monotonic decrease of the total mRNA counts with age during the early S phase, although it is consistent with evidence that transcription of a gene is downregulated before its replication occurs [95]. Using non-linear regression, the temporal variation of the mean count data in each of these four phases is found to be well fit by an exponential law *ke*^*βθ*^ where *k* and *β* are gene- and phase-specific constants (top right corner and bottom of Figure 3C).

In Materials and methods, we argue that the age-dependent mean mRNA predicted by our model will follow the empirical piecewise exponential law if the burst frequency or the mean burst size are chosen to vary exponentially with cell age. We make the following specific choices for the burst parameters. The burst frequency is assumed to change with cell age in a piecewise manner, from a constant *f*_1_ (for *θ < θ*_r_) to another constant *f*_2_ (for *θ* ≥ *θ*_r_). The mean burst size is assumed to be given by *b* = *ρe*^*βθ*^ where *ρ* and *β* are piecewise constant functions of *θ*. The functions specifying the variation of the burst frequency and the mean burst size with *θ* are illustrated on the right-hand side of Figure 3B.

Our functional choice for the burst parameters is in line with the findings from smFISH studies. These studies find that the burst frequency decreases upon DNA replication [20, 21] but does not change with cell volume, whereas the burst size increases with cell volume [21]. Given that cell age can be interpreted as a rough proxy for cell volume, these findings suggest that the burst size should depend on the cell age. Specifically, for exponentially growing cells that perfectly halve in size upon division and for which the growth rate is constant throughout the cell cycle, it is straightforward to show that *V* (*θ*) = *V*_0_*e*^log(2)*θ*^, where *V* (*θ*) is the cell volume at age *θ* and *V*_0_ is the cell birth volume. Since the burst size increases with cell volume, we choose the burst size to be *b* = *ρe*^*βθ*^, where we allow *ρ* and *β* to be some constants that are cell-cycle phase-dependent. This choice increases the flexibility of the model to fit the data and allows it to accommodate deviations due to incorrect or rough assumptions (such as constant growth rate). We note that the dependence of the burst size with cell age is a source of cell-to-cell variation within a cell-cycle phase, since there is a wide distribution of cell age within each phase. Hence, the age-dependent model overcomes the two limitations of the age-independent models mentioned earlier.

The equations for the temporal variation of the mean ⟨*m*⟩ (*θ*) and variance of mRNA counts *σ*^2^(*θ*) of the age-dependent model can be solved exactly using a computer algebra system (see Materials and methods). The distribution of mRNA counts at normalised time *θ* is then assumed to be given by a negative binomial distribution with mean ⟨*m*⟩ (*θ*) and variance *σ*^2^(*θ*); note that while the exact time-dependent distribution is not a negative binomial, this distribution is known to provide an excellent approximation for similar complex models of gene expression [69]. This choice is also motivated by the fact that the negative binomial model was the most commonly selected model when the scRNA-seq data was fit by the age-independent model (Materials and methods).

Finally, we use the method of maximum likelihood, with the likelihood being the aforementioned *θ*-dependent negative binomial, to fit the model to cell-age resolved data and estimate the parameters *f*_1_, *f*_2_, *ρ*_1_, *ρ*_2_, *β*_1_, *β*_2_, *β*_3_, *β*_4_ for each gene. In Figure 3D we show that this procedure leads to a good model fit to the data. Specifically, we show that for three sample genes the model predictions of the temporal dependence of the mean, variance and distribution of mRNA counts (obtained by evaluating the model using the estimated parameters) are in good agreement with the same statistics computed from the data. The accuracy of the inference procedure is further validated using simulated scRNA-seq data where the ground truth parameters are known (Supplementary Text S2; Supplementary Figure S2).

### Refinement of burst parameter inference using the cell-age dependent model

It is worthwhile to compare the burst parameters for each gene using the age-independent and the age-dependent models, since in this way we can determine the impact of the two main assumptions of the former age-independent model on the inference results. Because the two models differ in their parameterisation, a meaningful comparison must be done carefully, as follows.

Inference using the age-independent model led to a point estimate for the normalised burst frequency and normalised burst size in G1 and G2/M; multiplying these by the experimentally measured gene-specific mRNA degradation rate [66] (where it is available) leads to the absolute values of the burst parameters in the G1 and G2/M phases. In the age-dependent model, the burst frequency in G1 and G2/M phases simply corresponds to the parameters *f*_1_ and *f*_2_, respectively. To calculate the mean burst size for a gene in the G1 (G2/M) phase, we average the age-dependent burst size 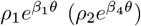 over the cell age distribution in this phase. For more details see Materials and methods. In Figure 4A, we contrast the phase-dependent burst parameters computed using the two models for 5 genes. Note that for these genes, the burst frequency estimated from the age-dependent model is larger than from the age-independent model. In contrast, the burst sizes appear to be smaller, although it is difficult to ascertain because of the large errors in the parameter estimates. In Figure 4B we confirm that for the vast majority of bursty genes using an age-independent model leads to a systematic underestimation of the burst frequency in both G1 and G2/M. Regarding the burst size, the age-independent model systematically overestimates it in G2/M; in G1, the burst size can be underestimated or overestimated with little influence on average. These results are broadly in agreement with a recent moment-based study that showed that extrinsic noise in parameter rates leads to systematic biases if they are inferred using a steady-state model that accounts only for intrinsic noise within each cell-cycle phase [19].

A main result of inference using the age-independent model was that the burst frequency decreases while the burst size increases as the cell cycle progresses from G1 to G2/M (Figure 2D). Specifically, we inferred that the genome-wide distribution of the ratio of burst frequency in G1 and G2/M for 1760 bursty genes has a median of 0.49 (1st and 3rd quartiles are 0.44 and 0.56), while for the burst size the median is 1.41 (1st and 3rd quartiles are 1.23 and 1.60). In contrast, the age-dependent model (for the subset of 1044 genes which can be described by this model; see Materials and methods) predicts that the burst frequency decreases, while the burst size remains approximately the same as the cell cycle progresses from G1 to G2/M (Figure 4C); the burst frequency and size ratios in G1 an G2/M have a median of 0.62 (1st and 3rd quartiles are 0.43 and 0.90) and of 0.86 (1st and 3rd quartiles are 0.64 and 1.15), respectively. Hence, the main difference between the predictions of the two models lies in how the burst size changes with cell-cycle phase.

In Supplementary Text S3, we redo the parameter inference using an alternative age-dependent model that unlike the model in Figure 3B does not assume a fixed cell age at which genes replicate in the S phase. The fits to the mean, variance and the distributions of mRNA counts as a function of the cell age of the two age-dependent models are of comparable quality, albeit the alternative model gives larger confidence intervals (data not shown). The burst frequency and size ratios between the G2/M and G1 phases estimated using the alternative age-dependent model are in good agreement with the original model estimates (Supplementary Figure S3), leading to the same conclusion as earlier, namely that the burst frequency decreases while the burst size remains approximately the same as the cell cycle progresses from G1 to G2/M. Specifically, the alternative age-dependent model (with age-dependent burst size and variable replication timing) estimates that the burst frequency and size ratios in G1 an G2/M have a median of 0.56 (1st and 3rd quartiles are 0.36 and 0.90) and of 1.05 (1st and 3rd quartiles are 0.71 and 1.57), respectively. In Supplementary Text S4, we test yet another alternative age-dependent model, where we now enforce an exponential scaling of transcription with cell age through an age-dependent burst frequency (and piecewise constant burst size), in contrast to the original model (Figure 3B) with an age-dependent burst size (and piecewise constant burst frequency). As shown in Supplementary Figure S4, this model formulation results in largely similar fits to the moments and distributions of the mRNA counts as the original model, and the estimates of burst parameter ratios between the G2/M and G1 phases are also in close agreement. Specifically, the alternative age-dependent model (with age-dependent burst frequency and fixed replication timing) estimates that the burst frequency and size ratios in G1 an G2/M have a median of 0.50 (1st and 3rd quartiles are 0.36 and 0.69) and of 1.09 (1st and 3rd quartiles are 0.86 and 1.31), respectively. These findings suggest that the choice of placement of an exponential scaling with cell age either on the burst size or burst frequency has little influence on the results of our transcriptome-wide analysis. Nevertheless, we choose to focus on the model with an age-dependent burst size throughout the paper, as such parametrisation has grounding in previous work (discussed in the last section).

We note that inference using both age-independent and age-dependent models leads to a negative correlation between the burst frequency and burst size ratios (this is clear from the downward-right trend of the grey scatter points in Figure 2D and Figure 4C). However, for the 1044 genes used in the age-dependent model inference, the correlation coefficient between the two ratios is −0.51 (the two alternative age-dependent models presented above have very similar associated correlation coefficients), while using the age-independent model for the same set of gene leads to a correlation coefficient of −0.82. This suggests that while some of the anti-correlation is an artefact of the assumptions behind the model used for inference, the rest may be reflecting underlying transcriptional mechanisms that co-regulate the burst parameters.

In the parameter estimation using age-independent models, technical noise was to some extent taken into account by fitting models that allow for zero-inflation. However, for the vast majority of genes, the zero-inflated models were rarely chosen by the model selection algorithm (Materials and methods). Whilst this implies that zero-inflation is likely not an issue for the 10x sequencing dataset we have analysed (in agreement with Ref. [51]), technical noise introduced by the inherent downsampling of mRNA counts may still be affecting the presented analysis. For example, the real transcript count for a particular gene in a cell might be 10 molecules but only 6 of these are captured — in this case there is no inflation of the zero count, but clearly technical noise has impacted the number of measured counts. This implies that downsampling could have possibly introduced a systematic bias in our estimation of the ratio of burst parameters in the two cell-cycle phases.

To address this concern, using a binomial mRNA capture model [84, 85] (illustrated in Figure 4D left), we developed a stochastic simulation-based strategy to estimate the bias introduced by ignoring this important source of technical noise in sequencing data. The idea is to simulate the true mRNA count data and then downsample it to resemble a virtual scRNA-seq experiment, fit the age-dependent model to the downsampled data and finally compare the estimated ratios of the burst parameters in G1 and G2/M (*Q*_*f*_ and *Q*_*b*_) to those obtained with the true count data. To mimic the true count data for a gene in a cell measured at a certain cell age, we sample the negative binomial distribution solution of the age-dependent model (Materials and methods). In this way, we generate data for 200 genes (each with a different set of rate parameters) in a population of cells, where the number of cells and the cell-age distribution match those of our mouse embryonic dataset. To mimic the observed scRNA-seq count data

*x* for a particular gene and cell, we downsample the true count *y* using the binomial model, i.e. *x* ∼ Binomial(*y, p*) where *p* is the capture probability for the cell which is assumed to be a random number drawn from the Beta distribution with mean ⟨*p*⟩ = 0.3 and coefficient of variation CV_*p*_ ≈ 0.15. In Figure 4D right we show a scatter plot of the burst parameter ratios between the G2/M and G1 phases estimated using the true count data and the corresponding downsampled (observed) data. Each point in the plot is a median over 100 repeats of the procedure, where the error bars indicate the lower and upper quartiles. Remarkably, our results show that downsampling due to a finite capture probability has very little effect on the estimated ratio of burst parameters in the two cell-cycle phases, and hence our inference results are robust with respect to technical noise.

## Discussion

In this paper, by fitting a variety of mechanistic models of stochastic gene expression to single-cell sequencing data that is cell-cycle specific, we have estimated the dependence of the burst parameters (the burst frequency and size) on the cell-cycle phase for about 1000 bursty genes in undifferentiated mouse embryonic stem cells. Inference using more sophisticated age-dependent models that account for noise due to transcriptional bursting, DNA replication, cell division and the coupling of gene expression with cell-cycle age reveals that, as the cell cycle progresses from G1 to G2/M, the median burst frequency decreases by about a half, while the median burst size remains approximately the same. However, the genome-wide distributions of the inferred ratios of burst frequency/size in G1 and G2/M are broad, thus suggesting a large degree of heterogeneity in transcriptional regulation patterns.

We note that whilst the burst parameters have been previously estimated using model-based inference applied to smFISH and scRNA-seq data, in the vast majority of cases, these were not estimated for individual cell-cycle phases [11–14, 19, 42, 44, 45, 48]. However, there are a few published studies which have reported this. An scRNA-seq-based study reported a comparison of single-allele burst parameters in G1, S and G2/M phases estimated using the telegraph model (see Extended Data Figure 10 in Ref [43]); because of the small numbers of cells in each phase (165, 28 and 31 in G1, S and G2/M, respectively) the errors in these parameters are necessarily large, and hence it is difficult to come to any reasonable conclusion on how the burst parameters vary across the cell cycle. Two smFISH-based studies [20, 22] used model-based inference to estimate the burst parameters before and after replication, but these were limited to only three eukaryotic genes and did not account for extrinsic noise due to factors varying within a cell-cycle phase, such as cell size. In Ref. [21] the burst parameter dependence with cell-cycle phase and cell size for about 25 mammalian genes was roughly estimated directly from the number of active transcription sites and their fluorescent intensity in smFISH measurements (without the use of model-based inference). Therefore, our study is the first to our knowledge that uses mechanistic model-based inference to reliably estimate the ratio of the burst parameters pre- and post-replication for over a thousand bursty eukaryotic genes.

Our results quantitatively agree with the findings of other studies and extend their predictions to a much larger number of genes. We found that partial dosage compensation is the norm across the transcriptome. This is because the distribution of the ratio of the mean counts in G2/M and G1 had a median close to 1.40, while perfect gene dosage compensation would be consistent with 1 and no dosage compensation with 2. This is in agreement with the findings of previous smFISH studies investigating gene expression in a few eukaryotic genes: in Ref. [22] it was found that for the Oct4 and Nanog genes, the fold change in the nascent mRNA level between the G1 and G2/M phases was 1.28 and 1.51 respectively, while in Ref. [20] for the GAL10 gene the fold change was measured to be 1.7. We inferred the transcriptome-wide distribution of *Q*_*f*_, the ratio of the burst frequencies per allele in G2/M and G1, and estimated its median to be equal to 0.56 or 0.60, depending on the choice of age-dependent model (referring to values given in Supplementary Text S3 both for the original model with age-dependent burst size with fixed replication timing and the alternative model with age-dependent burst size without fixed replication timing). This is in good agreement with the few cases in which these were estimated in previous smFISH studies of eukaryotic gene expression. In Ref. [22] for the Oct4 and Nanog genes in mouse embryonic stem cells, it was found that the burst frequency ratio was *Q*_*f*_ = 0.63 ± 0.06, while in Ref. [20] for the GAL10 gene in yeast, the frequency ratio was found to be *Q*_*f*_ = 0.66 ± 0.17. In Ref. [21] from smFISH measurements of the ratio of the number of active transcription sites in a cell and the total number of gene copies it was estimated that *Q*_*f*_ ≈ 0.5 for a few tens of genes in human primary fibroblast and lung cancer lines. We also found that the transcriptome-wide distribution of *Q*_*b*_, the ratio of the burst size per allele in G2/M and G1, has a median equal to 0.85 or 1.05 (reported in Supplementary Text S3 both for the original age-dependent model and the alternative model without fixed replication timing). This is in agreement with what was estimated in Ref. [21] for a few tens of mammalian genes.

Our study also sheds light on the relationship between the choice of model and the results of parameter inference. While we found that the values of the burst parameters were quite sensitive to the choice of the mechanistic model used for inference, the ratio of the burst frequency in the G1 and G2/M phases is a robust estimate. In contrast, the inference of the burst size ratio in G1 and G2/M is more sensitive to the chosen model and is best estimated by an age-dependent model, which accounts for cell-cycle dynamics and the coupling of the burst size to cell age. Surprisingly, we found that technical noise does not affect the estimation of the burst frequency and size ratios. We also found that the use of (age-independent) models that assume steady state within each cell-cycle phase and no extrinsic noise due to the coupling of transcription to cell age led to a strong artificial correlation between the ratio of the burst frequency in G1 and G2/M and the corresponding burst size ratio; models that do not make these assumptions (age-dependent models) find a smaller but sizeable correlation, thus suggesting that this may be due to underlying biological mechanisms.

In the future, the availability of high-quality sequencing data from a large population of cells of one type (tens of thousands) may make it possible to fit to the data more realistic models of gene expression than we have considered here. Examples of features that could be included are stochasticity in the duration of the cell-cycle [96], cell-size homeostasis mechanisms [97, 98] and multiple transcription and mRNA degradation steps [99], all of which have some impact on the mRNA count distribution and therefore could potentially lead to further refinements of the inferred burst parameter values.

In conclusion, we have devised a simple, tractable and robust approach to estimate from scRNA-seq data the dependence of the transcriptional burst parameters on the cell-cycle phase. While the use of mechanistic models for the extraction of the parameters controlling gene expression has become commonplace in fluorescence-based single-cell transcriptomics, their use in the field of single-cell sequencing has severely lagged behind [100]. This is largely due to the significant technical noise inherent to the sequencing technology and the difficulty of quantifying the sources of extrinsic noise. However, as we have here shown, with a carefully designed model-based inference approach, these issues present no impediment to building quantitative models of gene expression at the transcriptome level.

## Data Availability Statement

The original scRNA-seq dataset can be obtained from Ref. [57]. The processed data has been deposited at https://doi.org/10.5281/zenodo.10467234 and the code for this paper is available at https://github.com/augustinas1/CellCycle-RNAseq.

## Acknowledgments

This work was supported through an Alan Turing Institute Doctoral Studentship (EPSRC grant EP/N510129/1) and an ARC Laureate Fellowship (FL220100005) for A. S., and a Leverhulme Trust grant (Grant No. RPG-2020-327) for R. G. The authors would like to thank Kaan Öcal and Abhyudai Singh for useful discussions, as well as Michael Stumpf and the members of *Theoretical Systems Biology Group* at the University of Melbourne for their support. Lastly, the presented research was made possible by The University of Melbourne’s Research Computing Services and the Petascale Campus Initiative.

## Author contributions

Conceptualisation, R.G.; Methodology, A.S. and R.G.; Software, A.S.; Analysis, A.S.; Visualisation, A.S.; Writing – Original Draft, A.S. and R.G.; Writing – Review & Editing, A.S. and R.G.; Supervision, R.G.

## References

[1] A. Raj, C. S. Peskin, D. Tranchina, D. Y. Vargas, and S. Tyagi, “Stochastic mRNA synthesis in mammalian cells,” PLoS biology, 4(10):e309, 2006.

[2] D. M. Suter, N. Molina, D. Gatfield, K. Schneider, U. Schibler, and F. Naef, “Mammalian genes are transcribed with widely different bursting kinetics,” Science, 332(6028):472–474, 2011.

[3] B. T. Donovan, A. Huynh, D. A. Ball, H. P. Patel, M. G. Poirier, D. R. Larson, M. L. Ferguson, and T. L. Lenstra, “Live-cell imaging reveals the interplay between transcription factors, nucleosomes, and bursting,” The EMBO journal, 38(12):e100809, 2019.

[4] J. Rodriguez and D. R. Larson, “Transcription in living cells: Molecular mechanisms of bursting,” Annual review of biochemistry, 89:189–212, 2020.

[5] E. Tunnacliffe and J. R. Chubb, “What is a transcriptional burst?” Trends in Genetics, 36(4):288–297, 2020.

[6] A. Sanchez and I. Golding, “Genetic determinants and cellular constraints in noisy gene expression,” Science, 342(6163):1188–1193, 2013.

[7] I. Brouwer, H. P. Patel, J. V. W. Meeussen, W. Pomp, and T. L. Lenstra, “Single-molecule fluorescence imaging in living saccharomyces cerevisiae cells,” STAR protocols, 1(3):100142, 2020.

[8] T. L. Lenstra and D. R. Larson, “Single-molecule mRNA detection in live yeast,” Current protocols in molecular biology, 113(1):14–24, 2016.

[9] J. Peccoud and B. Ycart, “Markovian modeling of gene-product synthesis,” Theoretical population biology, 48(2):222– 234, 1995.

[10] J. Paulsson, O. G. Berg, and M. Ehrenberg, “Stochastic focusing: Fluctuation-enhanced sensitivity of intracellular regulation,” Proceedings of the National Academy of Sciences, 97(13):7148–7153, 2000.

[11] D. Zenklusen, D. R. Larson, and R. H. Singer, “Single-RNA counting reveals alternative modes of gene expression in yeast,” Nature structural & molecular biology, 15(12):1263–1271, 2008.

[12] A. Senecal, B. Munsky, F. Proux, N. Ly, F. E. Braye, C. Zimmer, F. Mueller, and X. Darzacq, “Transcription factors modulate c-Fos transcriptional bursts,” Cell reports, 8(1):75–83, 2014.

[13] H. Ochiai, T. Sugawara, T. Sakuma, and T. Yamamoto, “Stochastic promoter activation affects nanog expression variability in mouse embryonic stem cells,” Scientific reports, 4(1):7125, 2014.

[14] I. Golding, J. Paulsson, S. M. Zawilski, and E. C. Cox, “Real-time kinetics of gene activity in individual bacteria,” Cell, 123(6):1025–1036, 2005.

[15] N. Rosenfeld, J. W. Young, U. Alon, P. S. Swain, and M. B. Elowitz, “Gene regulation at the single-cell level,” Science, 307(5717):1962–1965, 2005.

[16] D. T. Gillespie, “Stochastic simulation of chemical kinetics,” Annual Review of Physical Chemistry, 58:35–55, 2007.

[17] M. B. Elowitz, A. J. Levine, E. D. Siggia, and P. S. Swain, “Stochastic gene expression in a single cell,” Science, 297(5584):1183–1186, 2002.

[18] J. M. Raser and E. K. O’Shea, “Control of stochasticity in eukaryotic gene expression,” Science, 304(5678):1811–1814, 2004.

[19] R. Grima and P.-M. Esmenjaud, “Quantifying and correcting bias in transcriptional parameter inference from single-cell data,” Biophysical Journal, 123:4–30, 2023.

[20] X. Fu, H. P. Patel, S. Coppola, L. Xu, Z. Cao, T. L. Lenstra, and R. Grima, “Quantifying how post-transcriptional noise and gene copy number variation bias transcriptional parameter inference from mRNA distributions,” Elife, 11:e82493, 2022.

[21] O. Padovan-Merhar, G. P. Nair, A. G. Biaesch, A. Mayer, S. Scarfone, S. W. Foley, A. R. Wu, L. S. Churchman, A. Singh, and A. Raj, “Single Mammalian Cells Compensate for Differences in Cellular Volume and DNA Copy Number through Independent Global Transcriptional Mechanisms,” Molecular Cell, 58(2):339–352, 2015.

[22] S. O. Skinner, H. Xu, S. Nagarkar-Jaiswal, P. R. Freire, T. P. Zwaka, and I. Golding, “Single-cell analysis of transcription kinetics across the cell cycle,” Elife, 5:e12175, 2016.

[23] X.-M. Sun, A. Bowman, M. Priestman, F. Bertaux, A. Martinez-Segura, W. Tang, C. Whilding, D. Dormann, V. Shahrezaei, and S. Marguerat, “Size-dependent increase in RNA polymerase II initiation rates mediates gene expression scaling with cell size,” Current Biology, 30(7):1217–1230, 2020.

[24] S. Berry and L. Pelkmans, “Mechanisms of cellular mRNA transcript homeostasis,” Trends in Cell Biology : 2022.

[25] I. G. Johnston, B. Gaal, R. P. d. Neves, T. Enver, F. J. Iborra, and N. S. Jones, “Mitochondrial variability as a source of extrinsic cellular noise,” PLoS computational biology, 8(3):e1002416, 2012.

[26] R. Foreman and R. Wollman, “Mammalian gene expression variability is explained by underlying cell state,” Molecular systems biology, 16(2):e9146, 2020.

[27] F. Tang, C. Barbacioru, Y. Wang, E. Nordman, C. Lee, N. Xu, X. Wang, J. Bodeau, B. B. Tuch, A. Siddiqui, et al., “mRNA-Seq whole-transcriptome analysis of a single cell,” Nature methods, 6(5):377–382, 2009.

[28] V. Svensson, R. Vento-Tormo, and S. A. Teichmann, “Exponential scaling of single-cell RNA-seq in the past decade,” Nature protocols, 13(4):599–604, 2018.

[29] G. X. Zheng, J. M. Terry, P. Belgrader, P. Ryvkin, Z. W. Bent, R. Wilson, S. B. Ziraldo, T. D. Wheeler, G. P. McDermott, J. Zhu, et al., “Massively parallel digital transcriptional profiling of single cells,” Nature communications, 8(1):14049, 2017.

[30] Z. Yao, H. Liu, F. Xie, S. Fischer, R. S. Adkins, A. I. Aldridge, S. A. Ament, A. Bartlett, M. M. Behrens, K. Van den Berge, et al., “A transcriptomic and epigenomic cell atlas of the mouse primary motor cortex,” Nature, 598(7879):103–110, 2021.

[31] S. Chen, P. Rivaud, J. H. Park, T. Tsou, E. Charles, J. R. Haliburton, F. Pichiorri, and M. Thomson, “Dissecting heterogeneous cell populations across drug and disease conditions with PopAlign,” Proceedings of the National Academy of Sciences, 117(46):28784–28794, 2020.

[32] S. Aibar, C.B. González-Blas, T. Moerman, V. A. Huynh-Thu, H. Imrichova, G. Hulselmans, F. Rambow, J.-C. Marine, P. Geurts, J. Aerts, et al., “SCENIC: single-cell regulatory network inference and clustering,” Nature methods, 14(11):1083–1086, 2017.

[33] T. Chari, G. Gorin, and L. Pachter, “Biophysically Interpretable Inference of Cell Types from Multimodal Sequencing Data,” bioRxiv : 2023–09, 2023.

[34] Q. Deng, D. Ramsköld, B. Reinius, and R. Sandberg, “Single-cell RNA-seq reveals dynamic, random monoallelic gene expression in mammalian cells,” Science, 343(6167):193–196, 2014.

[35] G. La Manno et al., “RNA velocity of single cells,” Nature, 560:494–498, 2018.

[36] V. Bergen, M. Lange, S. Peidli, F. A. Wolf, and F. J. Theis, “Generalizing RNA velocity to transient cell states through dynamical modeling,” Nature biotechnology, 38(12):1408–1414, 2020.

[37] D. A. Jaitin, E. Kenigsberg, H. Keren-Shaul, N. Elefant, F. Paul, I. Zaretsky, A. Mildner, N. Cohen, S. Jung, A. Tanay, et al., “Massively parallel single-cell RNA-seq for marker-free decomposition of tissues into cell types,” Science, 343(6172):776–779, 2014.

[38] A. P. Patel, I. Tirosh, J. J. Trombetta, A. K. Shalek, S. M. Gillespie, H. Wakimoto, D. P. Cahill, B. V. Nahed, W. T. Curry, R. L. Martuza, et al., “Single-cell RNA-seq highlights intratumoral heterogeneity in primary glioblastoma,” Science, 344(6190):1396–1401, 2014.

[39] C. Trapnell, D. Cacchiarelli, J. Grimsby, P. Pokharel, S. Li, M. Morse, N. J. Lennon, K. J. Livak, T. S. Mikkelsen, and J. L. Rinn, “The dynamics and regulators of cell fate decisions are revealed by pseudotemporal ordering of single cells,” Nature biotechnology, 32(4):381–386, 2014.

[40] B. Treutlein, D. G. Brownfield, A. R. Wu, N. F. Neff, G. L. Mantalas, F. H. Espinoza, T. J. Desai, M. A. Krasnow, and S. R. Quake, “Reconstructing lineage hierarchies of the distal lung epithelium using single-cell RNA-seq,” Nature, 509(7500):371–375, 2014.

[41] Z. Xue, K. Huang, C. Cai, L. Cai, C.-y. Jiang, Y. Feng, Z. Liu, Q. Zeng, L. Cheng, Y. E. Sun, et al., “Genetic programs in human and mouse early embryos revealed by single-cell RNA sequencing,” Nature, 500(7464):593–597, 2013.

[42] J. K. Kim and J. C. Marioni, “Inferring the kinetics of stochastic gene expression from single-cell RNA-sequencing data,” Genome biology, 14(1):1–12, 2013.

[43] A. J. M. Larsson, P. Johnsson, M. Hagemann-Jensen, L. Hartmanis, O. R. Faridani, B. Reinius, Å. Segerstolpe, C. M. Rivera, B. Ren, and R. Sandberg, “Genomic encoding of transcriptional burst kinetics,” Nature, 565:251–254, 2019.

[44] W. Tang, A. C. S. Jørgensen, S. Marguerat, P. Thomas, and V. Shahrezaei, “Modelling capture efficiency of single cell RNA-sequencing data improves inference of transcriptome-wide burst kinetics,” Bioinformatics, 39(7):btad395, 2023.

[45] K. Öcal, “Incorporating extrinsic noise into mechanistic modelling of single-cell transcriptomics,” bioRxiv : 2023–09, 2023.

[46] S. Luo, Z. Wang, Z. Zhang, T. Zhou, and J. Zhang, “Genome-wide inference reveals that feedback regulations constrain promoter-dependent transcriptional burst kinetics,” Nucleic Acids Research, 51(1):68–83, 2023.

[47] G. Gorin, J. J. Vastola, M. Fang, and L. Pachter, “Interpretable and tractable models of transcriptional noise for the rational design of single-molecule quantification experiments,” Nature Communications, 13(1):7620, 2022.

[48] Y. Jiang, N. R. Zhang, and M. Li, “SCALE: modeling allele-specific gene expression by single-cell RNA sequencing,” Genome biology, 18(1):1–15, 2017.

[49] J. K. Kim, A. A. Kolodziejczyk, T. Ilicic, S. A. Teichmann, and J. C. Marioni, “Characterizing noise structure in single-cell RNA-seq distinguishes genuine from technical stochastic allelic expression,” Nature communications, 6(1):8687, 2015.

[50] R. Jiang, T. Sun, D. Song, and J. J. Li, “Statistics or biology: the zero-inflation controversy about scRNA-seq data,” Genome Biology, 23(1):1–24, 2022.

[51] V. Svensson, “Droplet scrna-seq is not zero-inflated,” Nature Biotechnology, 38(2):147–150, 2020.

[52] Y. Cao, S. Kitanovski, R. Küppers, and D. Hoffmann, “UMI or not UMI, that is the question for scRNA-seq zero-inflation,” Nature Biotechnology, 39:158–159, 2021.

[53] T. Kivioja, A. Vähärautio, K. Karlsson, M. Bonke, M. Enge, S. Linnarsson, and J. Taipale, “Counting absolute numbers of molecules using unique molecular identifiers,” Nature methods, 9(1):72–74, 2012.

[54] C. Ziegenhain, B. Vieth, S. Parekh, B. Reinius, A. Guillaumet-Adkins, M. Smets, H. Leonhardt, H. Heyn, I. Hellmann, and W. Enard, “Comparative analysis of single-cell RNA sequencing methods,” Molecular cell, 65(4):631–643, 2017.

[55] D. Grün, L. Kester, and A. Van Oudenaarden, “Validation of noise models for single-cell transcriptomics,” Nature methods, 11(6):637–640, 2014.

[56] D. Volteras, V. Shahrezaei, and P. Thomas, “Global transcription regulation revealed from dynamical correlations in time-resolved single-cell RNA-sequencing,” bioRxiv : 2023–10, 2023.

[57] A. Riba, A. Oravecz, M. Durik, S. Jiménez, V. Alunni, M. Cerciat, M. Jung, C. Keime, W. M. Keyes, and N. Molina, “Cell cycle gene regulation dynamics revealed by RNA velocity and deep-learning,” Nature Communications, 13(2865):1–13, 2022.

[58] V. Bergen, R. A. Soldatov, P. V. Kharchenko, and F. J. Theis, “RNA velocity—current challenges and future perspectives,” Molecular Systems Biology, 17(8):e10282, 2021.

[59] G. Gorin, M. Fang, T. Chari, and L. Pachter, “RNA velocity unraveled,” PLOS Computational Biology, 18(9):e1010492, 2022.

[60] A. R. Stinchcombe, C. S. Peskin, and D. Tranchina, “Population density approach for discrete mRNA distributions in generalized switching models for stochastic gene expression,” Physical Review E, 85(6):061919, 2012.

[61] S. C. Hicks, F. W. Townes, M. Teng, and R. A. Irizarry, “Missing data and technical variability in single-cell RNA-sequencing experiments,” Biostatistics, 19(4):562–578, 2018.

[62] K. Choi, Y. Chen, D. A. Skelly, and G. A. Churchill, “Bayesian model selection reveals biological origins of zero inflation in single-cell transcriptomics,” Genome Biology, 21(1):1–16, 2020.

[63] W. Chen, Y. Li, J. Easton, D. Finkelstein, G. Wu, and X. Chen, “UMI-count modeling and differential expression analysis for single-cell RNA sequencing,” Genome Biology, 19(1):1–17, 2018.

[64] F. W. Townes, S. C. Hicks, M. J. Aryee, and R. A. Irizarry, “Feature selection and dimension reduction for single-cell RNA-Seq based on a multinomial model,” Genome Biology, 20(1):1–16, 2019.

[65] A. Waisman, F. Sevlever, M. Elías Costa, M. S. Cosentino, S. G. Miriuka, A. C. Ventura, and A. S. Guberman, “Cell cycle dynamics of mouse embryonic stem cells in the ground state and during transition to formative pluripotency,” Scientific Reports, 9(8051):1–10, 2019.

[66] L. V. Sharova, A. A. Sharov, T. Nedorezov, Y. Piao, N. Shaik, and M. S. Ko, “Database for mRNA half-life of 19 977 genes obtained by DNA microarray analysis of pluripotent and differentiating mouse embryonic stem cells,” DNA research, 16(1):45–58, 2009.

[67] I. Hiratani, T. Ryba, M. Itoh, T. Yokochi, M. Schwaiger, C.-W. Chang, Y. Lyou, T. M. Townes, D. Schübeler, and D. M. Gilbert, “Global reorganization of replication domains during embryonic stem cell differentiation,” PLoS biology, 6(10):e245, 2008.

[68] O. G. Berg, “A model for the statistical fluctuations of protein numbers in a microbial population,” Journal of theoretical biology, 71(4):587–603, 1978.

[69] Z. Cao and R. Grima, “Analytical distributions for detailed models of stochastic gene expression in eukaryotic cells,” Proceedings of the National Academy of Sciences, 117(9):4682–4692, 2020.

[70] J. Bezanson, A. Edelman, S. Karpinski, and V. B. Shah, “Julia: A Fresh Approach to Numerical Computing,” SIAM Review : 2017.

[71] M. Besançon, T. Papamarkou, D. Anthoff, A. Arslan, S. Byrne, D. Lin, and J. Pearson, “Distributions.jl: Definition and Modeling of Probability Distributions in the JuliaStats Ecosystem,” Journal of Statistical Software, 98(16):1–30, 2021.

[72] S. Danisch and J. Krumbiegel, “Makie.jl: Flexible high-performance data visualization for Julia,” Journal of Open Source Software, 6(65):3349, 2021.

[73] T. N. Vu, Q. F. Wills, K. R. Kalari, N. Niu, L. Wang, M. Rantalainen, and Y. Pawitan, “Beta-Poisson model for single-cell RNA-seq data analyses,” Bioinformatics, 32(14):2128–2135, 2016.

[74] R. Feldt and A. Stukalov, BlackBoxOptim.jl, 2018.

[75] P. K. Mogensen and A. N. Riseth, “Optim: A mathematical optimization package for Julia,” Journal of Open Source Software, 3(24):615, 2018.

[76] A. E. Raftery, “Bayesian Model Selection in Social Research,” Sociological Methodology, 25:111–163, 1995.

[77] A. A. Neath and J. E. Cavanaugh, “The Bayesian information criterion: background, derivation, and applications,” WIREs Computational Statistics, 4(2):199–203, 2012.

[78] T. O. Kvålseth, “Cautionary Note about R^2^,” The American Statistician, 39(4):279–285, 1985.

[79] Y. Pawitan, In All Likelihood. Oxford, England, UK: Oxford University Press, 2013.

[80] C. Kreutz, A. Raue, D. Kaschek, and J. Timmer, “Profile likelihood in systems biology,” FEBS Journal, 280(11):2564–2571, 2013.

[81] C. Kreutz, A. Raue, and J. Timmer, “Likelihood based observability analysis and confidence intervals for predictions of dynamic models,” BMC Systems Biology, 6(1):1–9, 2012.

[82] A. Wächter and L. T. Biegler, “On the implementation of an interior-point filter line-search algorithm for large-scale nonlinear programming,” Mathematical Programming, 106(1):25–57, 2006.

[83] B. Legat, O. Dowson, J. Dias Garcia, and M. Lubin, “MathOptInterface: A data structure for mathematical optimization problems,” INFORMS Journal on Computing, 34(2):672–689, 2021.

[84] A. M. Klein, L. Mazutis, I. Akartuna, N. Tallapragada, A. Veres, V. Li, L. Peshkin, D. A. Weitz, and M. W. Kirschner, “Droplet barcoding for single-cell transcriptomics applied to embryonic stem cells,” Cell, 161(5):1187–1201, 2015.

[85] W. Tang, F. Bertaux, P. Thomas, C. Stefanelli, M. Saint, S. Marguerat, and V. Shahrezaei, “BayNorm: Bayesian gene expression recovery, imputation and normalization for single-cell RNA-sequencing data,” Bioinformatics, 36(4):1174–1181, 2020.

[86] A. M. Klein, L. Mazutis, I. Akartuna, N. Tallapragada, A. Veres, V. Li, L. Peshkin, D. A. Weitz, and M. W. Kirschner, “Droplet Barcoding for Single-Cell Transcriptomics Applied to Embryonic Stem Cells,” Cell, 161(5):1187–1201, 2015.

[87] Y. Wan, D. G. Anastasakis, J. Rodriguez, M. Palangat, P. Gudla, G. Zaki, M. Tandon, G. Pegoraro, C. C. Chow, M. Hafner, et al., “Dynamic imaging of nascent rna reveals general principles of transcription dynamics and stochastic splice site selection,” Cell, 184(11):2878–2895, 2021.

[88] S. Luo, Z. Zhang, Z. Wang, X. Yang, X. Chen, T. Zhou, and J. Zhang, “Inferring transcriptional bursting kinetics from single-cell snapshot data using a generalized telegraph model,” Royal Society Open Science, 10(4):221057, 2023.

[89] Z. Cao, T. Filatova, D. A. Oyarzún, and R. Grima, “A stochastic model of gene expression with polymerase recruitment and pause release,” Biophysical Journal, 119(5):1002–1014, 2020.

[90] A. G. Nicoll, J. Szavits-Nossan, M. R. Evans, and R. Grima, “Transient power-law behaviour following induction distinguishes between competing models of stochastic gene expression,” bioRxiv : 2023–12, 2023.

[91] Y. Wang, J. Szavits-Nossan, Z. Cao, and R. Grima, “Joint distribution of nuclear and cytoplasmic mrna levels in stochastic models of gene expression: Analytical results and parameter inference,” bioRxiv : 2024–04, 2024.

[92] M. Allen, D. Poggiali, K. Whitaker, T. R. Marshall, J. van Langen, and R. A. Kievit, “Raincloud plots: a multi-platform tool for robust data visualization,” Wellcome Open Research, 4:63.: 2021. eprint: 31069261.

[93] M. R. Dowling, A. Kan, S. Heinzel, J. H. Zhou, J. M. Marchingo, C. J. Wellard, J. F. Markham, and P. D. Hodgkin, “Stretched cell cycle model for proliferating lymphocytes,” Proceedings of the National Academy of Sciences, 111(17):6377–6382, 2014.

[94] C. Jia and R. Grima, “Coupling gene expression dynamics to cell size dynamics and cell cycle events: Exact and approximate solutions of the extended telegraph model,” iScience, 26(1):105746, 2023.

[95] I. Tsirkas, D. Dovrat, M. Thangaraj, I. Brouwer, A. Cohen, Z. Paleiov, M. M. Meijler, T. Lenstra, and A. Aharoni, “Transcription-replication coordination revealed in single live cells,” Nucleic Acids Research, 50(4):2143–2156, 2022.

[96] R. Perez-Carrasco, C. Beentjes, and R. Grima, “Effects of cell cycle variability on lineage and population measurements of messenger rna abundance,” Journal of the Royal Society Interface, 17(168):20200360, 2020.

[97] C. Jia and R. Grima, “Frequency domain analysis of fluctuations of mrna and protein copy numbers within a cell lineage: Theory and experimental validation,” Physical Review X, 11(2):021032, 2021.

[98] C. Jia, A. Singh, and R. Grima, “Concentration fluctuations in growing and dividing cells: Insights into the emergence of concentration homeostasis,” PLoS Computational Biology, 18(10):e1010574, 2022.

[99] D. E. Weidemann, J. Holehouse, A. Singh, R. Grima, and S. Hauf, “The minimal intrinsic stochasticity of constitutively expressed eukaryotic genes is sub-Poissonian,” Science Advances, 9(32):eadh5138, 2023.

[100] G. Gorin, J. J. Vastola, and L. Pachter, “Studying stochastic systems biology of the cell with single-cell genomics data,” Cell Systems, 14(10):822–843, 2023.

