## Supplementary Information for "Cell-cycle dependence of bursty gene expression: insights from fitting mechanistic models to single-cell RNA-seq data"

### Supplementary Text S1: Validation of steady-state model selection

In this section, we conduct a small benchmark study to assess the robustness of our steady-state model selection algorithm based on the Bayesian Information Criterion (BIC), described in Methods. Note that although the BIC is a stringent test only in the asymptotic limit (assuming the dataset is large enough), we expect it to be suitable in our case due to fairly large sample sizes (941 cells in the G1 phase and 2113 cells in the G2/M phase). Furthermore, its strong penalisation for models with a larger number of parameters (its preference for the most parsimonious model) is also practically very useful in reducing the inherent model unidentifiability and hence minimising the errors in parameter estimates.

We outline the procedure testing the model selection for Poisson-like model fits as follows. We select a random subset of 500 genes out of all genes that were found to be optimally fit by the Poisson model in the G1 phase. For each gene, we sample the corresponding Poisson distribution $941 \times 2$ times to construct a dataset equivalent in structure to the original mRNA count data of all cells in the G1 phase (the factor 2 accounts for the fact that we have two independent gene copies). We repeat the procedure 10 times to generate 10 sampled count datasets, and rerun the inference and model selection procedure using all six steady-state models for the 10 resampled count datasets — for each gene, we pick the model that was most commonly selected as the best-fit model out of 10 re-runs of the procedure.

In Figure S1A left, the corresponding optimal model fits are visualised as a bar plot, showing what fraction of the selected 500 Poisson-like genes were found to be best-fit by different steady-state models. In this case, the Poisson model is preferred in all cases, which is expected given that the BIC usually favours the most parsimonious model. Furthermore, we repeat the same procedure for random subsets of 100 genes that were found to be best-fit by the zero-inflated Poisson (Figure S1A middle) or negative binomial (Figure S1A right) models, showing that for the majority of genes the data-generating model was selected as the optimal model, and hence validating our model selection procedure.

Note that we cannot perform the same test for the more complicated zero-inflated negative binomial, telegraph and zero-inflated telegraph models, because we find only a handful of genes that are optimally fit by these models (as noted in Methods). Although we could randomly sample the parameter space to generate the models and redo the benchmarking in this fashion, the results would be harder to interpret due to the inherent nesting of the presented steady-state models. For example, as discussed in Methods, the steady-state solution of the telegraph model converges to a Poisson distribution in the constitutive limit and to a negative binomial distribution in the bursty limit — by randomly parametrising the telegraph model and redoing the inference we would often find a Poisson or a negative binomial as the optimal model.

We illustrate this point in Figure S1B, where we perform a similar inference and model selection validation study by generating synthetic data using the telegraph model for different values of the switching on rate $\sigma_1$ (gene switching from the transcriptionally inactive state to the active state), while keeping other parameters fixed as $\sigma_0 = 1$ (gene switching off rate), $\rho = 50$ (mRNA production rate) and $d = 1$ (degradation rate). Here we consider 100 different values of $\sigma_1$ evenly spaced in the log10 space from $10^{-2}$ to $10^2$ — note that the combination of the gene switching rates can also be interpreted in terms of the time spent in the transcriptionally active state, given by $f_{\text{on}} = \frac{\sigma_1}{\sigma_0+\sigma_1}$ (superimposed in Figure S1B as a solid purple line).

For each value of $\sigma_1$, we resample the corresponding telegraph model solution to construct a dataset equivalent in structure to the original count data in the G1 phase (as done previously in Fig S1A). We resample 100 similar datasets for each $\sigma_1$, which we then fit to all six age-independent steady-state models — in Figure S1B we plot the frequency at which each model was selected as the optimal one. Note that the zero-inflated telegraph model is not colour-coded because it was never picked as the optimal model. As expected from the theoretical analysis, our model selection algorithm picks the Poisson (negative binomial) model as the optimal model in the constitutive (bursty) limit, which corresponds to a high (low) value of $f_{\text{on}}$. In these limits, where the telegraph model converges to simpler distributions, its individual parameter values become unidentifiable, and hence choosing the corresponding simpler model is preferable in order to reduce the estimation errors, which our model selection procedure successfully achieves as demonstrated.

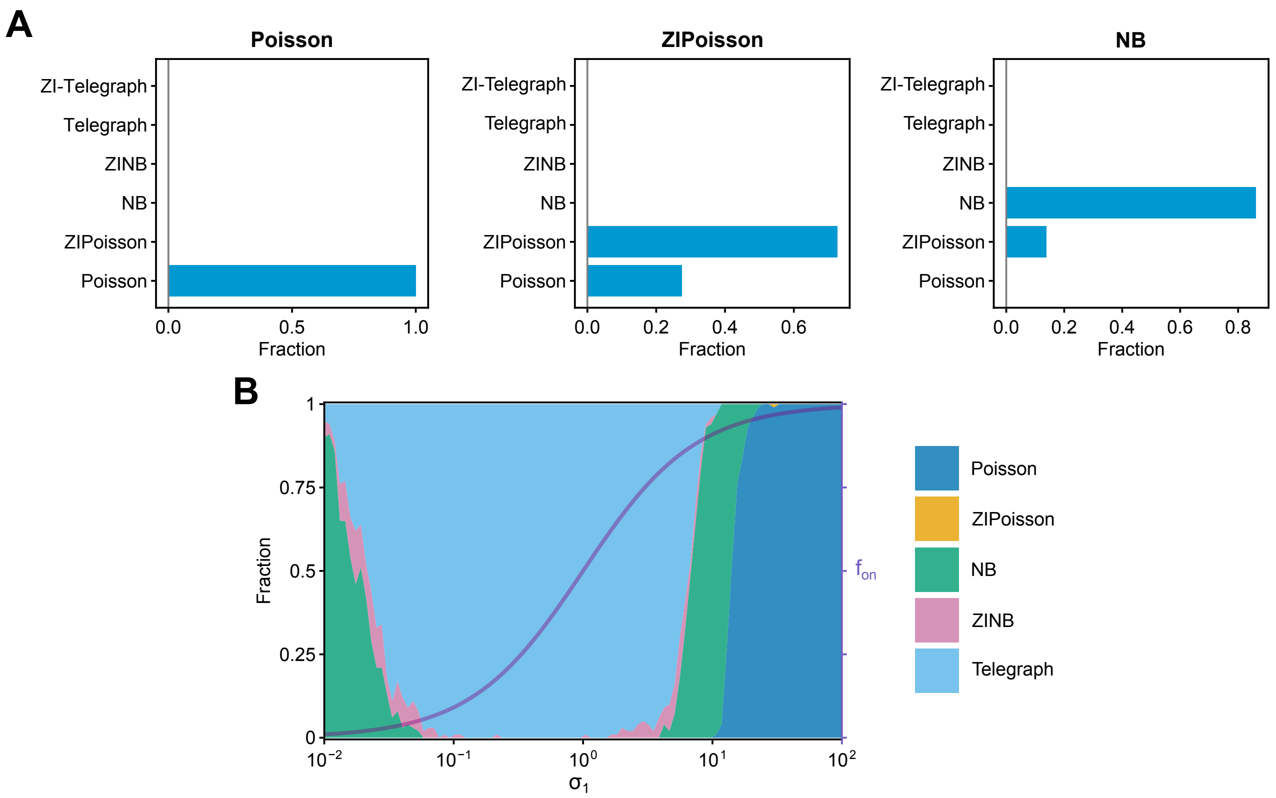


**Figure S1:** Verification of the steady-state model selection procedure. **(A)** Each bar plot shows the fraction of times each of the six steady-state age-independent models was selected, whereas the plot title indicates the model used to generate the data (that best fits the original count data). We perform the test for Poisson (left), zero-inflated Poisson (ZIPoisson) (middle) and negative binomial (NB) (right) models. **(B)** Fraction of times at which each steady-state model is selected, where the data is generated from the telegraph model at different switching on rates $\sigma_1$ (with other parameters kept fixed). The purple solid line indicates the corresponding $f_{\mathrm{on}}$, i.e. fraction of time spent by the gene in the transcriptionally active state. See Supplementary Text S1 for more details.

### Supplementary Text S2: Validation of age-dependent model inference

To ensure that our age-dependent model parameter estimates are accurate, we perform a self-consistency check using synthetically generated data. We consider 200 randomly selected genes from the final set of 1044 genes that are found to be well-characterised by the age-dependent transcriptional model described in the main text. For each gene, we use the model with the parameters best-fit to the experimental mRNA count data to construct a corresponding synthetic dataset — by fitting this data and comparing the obtained parameter estimates to their ground truth we can probe the validity of our inference strategy.

To generate synthetic count data for each gene, we use the true model parameters $\{f_1, f_2, \rho_1, \rho_2, \beta_1, \beta_2, \beta_3, \beta_4\}$ to build and parametrise the corresponding chemical reaction network in a piecewise fashion (as discussed in Methods and visualised in Figure 3B). Instead of sampling the approximate negative binomial solution of the chemical master equation describing the model, we simulate it exactly using the Stochastic Simulation Algorithm (SSA) [1]. This is done in Julia using Catalyst.jl [2] for model implementation and DifferentialEquations.jl [3] for stochastic simulation.

Our simulation setup corresponds to a cell lineage experiment: after the division of a mother cell, we randomly pick a single daughter cell and continue tracking its progression throughout the cell cycle. Starting from the initial condition of zero mRNA molecules, we simulate the model until time $t = 10T$, corresponding to 10 full cell-cycle durations, where we again use the experimentally determined median cell-cycle duration $T = 13.25\,\text{h}$ for mESCs [4] and the gene-specific degradation rates [5]. Running the simulation for 10 complete cell cycles is a heuristically motivated choice, as we expect the system at this time to have reached complete cyclo-stationarity (or steady-state growth), so that the mRNA distribution at cell age $\theta$ does not depend on which generation the cell belongs to. By visual inspection, cyclo-stationary values of the means and variances are usually achieved as early as by the third iteration (data not shown).

We create a synthetic dataset for a single gene by sampling 5294 SSA realisations (equivalent to the number of cells in the experimental dataset, as noted in Methods), and considering the mRNA counts at simulation times $t \in [9T, 10T)$, corresponding to the 10th cell-cycle generation. By recording the mRNA counts of each SSA trajectory at certain values of $\theta$, we obtain a dataset analogous in structure to the experimental count data, and use it to fit the age-dependent model.

We repeat the data generation and inference procedure 100 times for each gene and report the median observed parameter estimates and their interquartile ranges in Figure S2. In most cases, the observed parameters are close to their true values, with the exception of the parameter $\beta_2$ that displays a relatively large level of variability. Note that $\beta_2$ is usually negative and for the majority of genes tends to the value of -100 (the artificial parameter bound used to constrain the search space), making it quite sensitive to noise in the data and leading to a higher degree of parameter unidentifiability. Nevertheless, the observed estimates of $\beta_2$ capture the downward trend of mRNA counts in the early S phase, and crucially, the predicted burst frequency and burst size ratios between the G2/M and G1 phases remain accurate (bottom row of Figure S2).

Overall, the observed parameter estimates agree with the ground-truth values and confirm the robustness of our inference method used to fit the experimental data. Moreover, the results of this analysis validate the model assumption made in Methods, where we approximate the true mRNA count distribution at cell age $\theta$ by a negative binomial distribution parametrised by the exact moments of the age-dependent model evaluated at $\theta$.

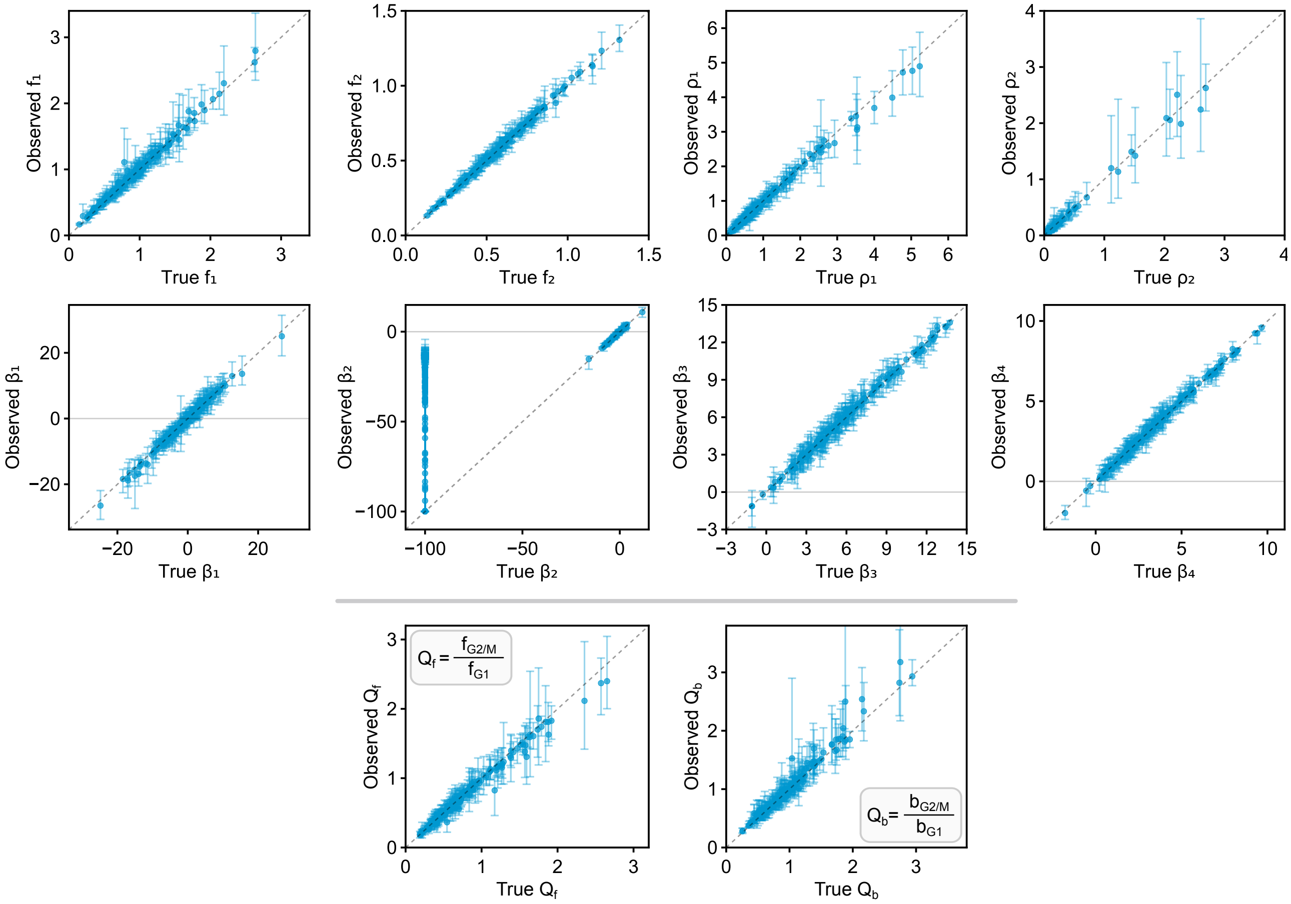


**Figure S2:** Validation of the model fitting procedure for the age-dependent model using synthetic scRNA-seq data. Scatter plots show the observed fits of each age-dependent model parameter and the estimated burst frequency and burst size ratios between the G2/M and G1 phases, compared to their ground-truth values. Each data point is the median over 100 repeats of the synthetic data generation and inference procedure, where the error bars denote the lower and upper quartiles — see Supplementary Text S2 for more details.

#### Supplementary Text S3: Alternative model with age-dependent burst size & variable replication timing

In this section, we consider an alternative form of the age-dependent bursty model of gene expression which differs from the model presented in Methods only by its piecewise parameterisation in the S phase. Namely, instead of defining allele-specific parameters prior- and post-replication, we take a more general approach and consider the total burst frequency and the total mean burst size over all gene copies in the S phase. This allows us to avoid the rough assumption that DNA replication occurs in the middle of the S phase at cell age $\theta_r$ for all genes. In what follows, we derive an approximate solution to the alternative model in terms of the age-dependent count distribution which we use to perform transcriptome-wide inference, and finally compare the model predictions to those of the age-dependent model utilised in the main text.

Here we assume the piecewise parameters $f$ and $\rho$ change from constants $f_1$ and $\rho_1$ in the G1 phase for $\theta < \theta_{\text{G1}}$ (per-allele values assuming 2 gene copies) to $f_2$ and $\rho_2$ in the S phase for $\theta_{\text{G1}} \leq \theta < \theta_{\text{S}}$ (total values over all alleles), and finally to $f_3$ and $\rho_3$ in the G2/M phase for $\theta \geq \theta_{\text{S}}$ (per-allele values assuming 4 gene copies). We again split the parameter $\beta$ into four parts for extra model flexibility required to fit the non-monotonic variation of the mean mRNA counts with cell age in the S phase. However, we now split the S phase into early and late stages at $\theta_{\min} = 0.33$, which denotes the $\theta$ bin with the minimum total mRNA count per cell observed in the S phase, as highlighted in Figure S3A. Note that $\theta_{\min}$ does not coincide with the assumed replication time $\theta_r = 0.39$ used to parametrise the age-dependent model in the main text. The parameter $\beta$ is given by the piecewise constant $\beta_1$ in the G1 phase for $\theta < \theta_{\text{G1}}$, $\beta_2$ in the early S phase for $\theta_{\text{G1}} \leq \theta < \theta_{\min}$, $\beta_3$ in the late S phase for $\theta_{\min} \leq \theta < \theta_{\text{S}}$ and $\beta_4$ in the G2/M phase for $\theta \geq \theta_{\text{S}}$. A schematic illustration of the model and the piecewise changes in model parameters with cell age $\theta$ are illustrated in Figure S3B.

As done previously, using the chemical master equation for the reaction scheme of the bursty model (Eq. (9) in Methods), we derive piecewise differential equations for the mean mRNA counts due to all alleles in each cell-cycle phase:

$$\partial_\theta \langle m\rangle_1(\theta) = 2f_1\rho_1 T e^{\beta_1\theta} - dT\langle m\rangle_1(\theta), \quad \theta \in [0, \theta_{\text{G1}}], \tag{1}$$
$$\partial_\theta \langle m\rangle_2(\theta) = f_2\rho_2 T e^{\beta_2\theta} - dT\langle m\rangle_2(\theta), \quad \theta \in [\theta_{\text{G1}}, \theta_{\min}] \tag{2}$$
$$\partial_\theta \langle m\rangle_3(\theta) = f_2\rho_2 T e^{\beta_3\theta} - dT\langle m\rangle_3(\theta), \quad \theta \in [\theta_{\min}, \theta_{\text{S}}], \tag{3}$$
$$\partial_\theta \langle m\rangle_4(\theta) = 4f_3\rho_3 T e^{\beta_4\theta} - dT\langle m\rangle_4(\theta), \quad \theta \in [\theta_{\text{S}}, 1], \tag{4}$$

where the factors of 2 and 4 correspond to the number of alleles in the G1 and G2/M phases, whereas in the S phase we absorb the unknown number of alleles in the total burst frequency $f_2$. The four boundary conditions are equivalent in form to Eqs. (17)-(20) in Methods:

$$\langle m\rangle_1(\theta_{\text{G1}}) = \langle m\rangle_2(\theta_{\text{G1}}), \tag{5}$$
$$\langle m\rangle_2(\theta_{\min}) = \langle m\rangle_3(\theta_{\min}), \tag{6}$$
$$\langle m\rangle_3(\theta_{\text{S}}) = \langle m\rangle_4(\theta_{\text{S}}), \tag{7}$$
$$\langle m\rangle_1(0) = \frac{1}{2}\langle m\rangle_4(1). \tag{8}$$

Similarly, the piecewise differential equations for the variances of mRNA counts are given by

$$\partial_\theta\sigma_1^2(\theta) = -2dT\sigma_1^2(\theta) + 2f_1\rho_1 T e^{\beta_1\theta}\left(1 + 2\rho_1 e^{\beta_1\theta}\right) + dT\langle m\rangle_1(\theta), \quad \theta \in [0, \theta_{\text{G1}}], \tag{9}$$
$$\partial_\theta\sigma_2^2(\theta) = -2dT\sigma_2^2(\theta) + f_2\rho_2 T e^{\beta_2\theta}\left(1 + 2\rho_2 e^{\beta_2\theta}\right) + dT\langle m\rangle_2(\theta), \quad \theta \in [\theta_{\text{G1}}, \theta_{\min}], \tag{10}$$
$$\partial_\theta\sigma_3^2(\theta) = -2dT\sigma_3^2(\theta) + f_2\rho_2 T e^{\beta_3\theta}\left(1 + 2\rho_2 e^{\beta_3\theta}\right) + dT\langle m\rangle_3(\theta), \quad \theta \in [\theta_{\min}, \theta_{\text{S}}], \tag{11}$$
$$\partial_\theta\sigma_4^2(\theta) = -2dT\sigma_4^2(\theta) + 4f_3\rho_3 T e^{\beta_4\theta}\left(1 + 2\rho_3 e^{\beta_4\theta}\right) + dT\langle m\rangle_4(\theta), \quad \theta \in [\theta_{\text{S}}, 1], \tag{12}$$

and the associated boundary conditions are

$$\sigma_1^2(\theta_{\text{G1}}) = \sigma_2^2(\theta_{\text{G1}}), \tag{13}$$
$$\sigma_2^2(\theta_{\min}) = \sigma_3^2(\theta_{\min}), \tag{14}$$
$$\sigma_3^2(\theta_{\text{S}}) = \sigma_4^2(\theta_{\text{S}}), \tag{15}$$
$$\sigma_1^2(0) = \frac{\sigma_4^2(1) + \langle m\rangle_4(1)}{4}. \tag{16}$$

As before, the complete ODE system can be solved using Mathematica, and we approximate the true mRNA count distribution at any $\theta$ using a negative binomial parametrised by the age-dependent moment solutions:

$$P(m,\theta) = \mathrm{NB}\left(m; \frac{(\langle m\rangle_1(\theta))^2}{\sigma_1^2(\theta) - \langle m\rangle_1(\theta)}, \frac{\langle m\rangle_1(\theta)}{\sigma_1^2(\theta)}\right), \quad \theta \in [0, \theta_{\mathrm{G1}}], \tag{17}$$

$$P(m,\theta) = \mathrm{NB}\left(m; \frac{(\langle m\rangle_2(\theta))^2}{\sigma_2^2(\theta) - \langle m\rangle_2(\theta)}, \frac{\langle m\rangle_2(\theta)}{\sigma_2^2(\theta)}\right), \quad \theta \in [\theta_{\mathrm{G1}}, \theta_{\min}], \tag{18}$$

$$P(m,\theta) = \mathrm{NB}\left(m; \frac{(\langle m\rangle_3(\theta))^2}{\sigma_3^2(\theta) - \langle m\rangle_3(\theta)}, \frac{\langle m\rangle_3(\theta)}{\sigma_3^2(\theta)}\right), \quad \theta \in [\theta_{\min}, \theta_{\mathrm{S}}], \tag{19}$$

$$P(m,\theta) = \mathrm{NB}\left(m; \frac{(\langle m\rangle_4(\theta))^2}{\sigma_4^2(\theta) - \langle m\rangle_4(\theta)}, \frac{\langle m\rangle_4(\theta)}{\sigma_4^2(\theta)}\right), \quad \theta \in [\theta_{\mathrm{S}}, 1]. \tag{20}$$

We fit the obtained age-dependent model to $\theta$-resolved mRNA count data using the method of maximum likelihood and estimate the parameters $f_1, f_2, f_3, \rho_1, \rho_2, \rho_3, \beta_1, \beta_2, \beta_3, \beta_4$ for each of the 1351 bursty genes (obtained after the initial gene filtering steps outlined in Methods. Note that the employed inference procedure and the parameter constraints are equivalent to those described for the original age-dependent model in Methods. We next filter out genes with poor model fits that predict a negative correlation of the mean counts with cell age in the G1 or G2/M phases, or have a negative associated $R^2$ value for the predicted mean and/or variance throughout the full cell cycle, as done in Methods. In addition, we discard a handful of genes that in any cell-cycle phase have an associated burst frequency $f$ tending to the upper parameter bound, i.e. enforcing $f_1, f_2, f_3 < 100$, because higher $f$ values are indicative of a Poisson-like fit that can lead to physically unrealistic mean burst size and parameter ratio estimates. As a final step, we also remove a few genes that have an estimated burst frequency or burst size ratio between the G2/M and G1 phases greater than 10 — such values indicate extreme outliers in the dataset, in part caused by the larger errors on the parameter estimates due to additional complexity of the alternative age-dependent model, and which may heavily skew the computed statistics. After these filtering steps, we are left with 1063 genes, 958 of which intersect with the 1044 genes obtained using the age-dependent model in the main text. Therefore, for the model comparison below, we consider the overlapping set of 958 genes and their model fits.

In Figure S3C we compare the inferred burst frequencies and the mean burst sizes between the two age-dependent models in both G1 and G2/M phases for all 958 overlapping genes. Although there is an observable spread in the burst parameters, on the median-level they remain largely similar, with a larger discrepancy observed only for the mean burst sizes in the G1 phase. Nevertheless, in Figure S3D we confirm that the alternative age-dependent model estimates of burst frequency and size ratios between the G2/M and G1 phases are qualitatively similar to the same estimates computed with the age-dependent model from the main text. More specifically, the alternative age-dependent model predicts that the burst frequency and size ratios between G2/M and G1 have a median of 0.56 (1st and 3rd quartiles are 0.36 and 0.90) and of 1.05 (1st and 3rd quartiles are 0.71 and 1.57), respectively, whereas for the main-text model the median burst frequency ratio is 0.60 (1st and 3rd quartiles are 0.42 and 0.87) and the median burst size ratio is 0.85 (1st and 3rd quartiles are 0.64 and 1.13). Note that the quantiles for the main-text model here are slightly different to those reported in Results, as we now consider only 958 genes. Overall, these findings agree with the results in the main text, similarly suggesting that the burst frequency decreases and the burst size stays approximately the same as the cells progress from G1 to G2/M. We also observe that both age-dependent models lead to a similar negative correlation between the burst frequency and burst size ratios: for the set of 958 genes the correlation coefficient is -0.50 using the alternative age-dependent model, and -0.55 using the main-text model (the correlations are weaker than observed for the age-independent models, as discussed in Results).

The consistent predictions of the two models bring more confidence that the approximation of a fixed DNA replication time in the middle of the S phase for the age-dependent model in the main text does not significantly affect the results of the transcriptome-wide analysis of burst parameters. Finally, we note that due to fewer parameters, inference for the age-dependent model in the main text leads to tighter confidence intervals and hence less parameter unidentifiability than for the alternative model (data not shown).

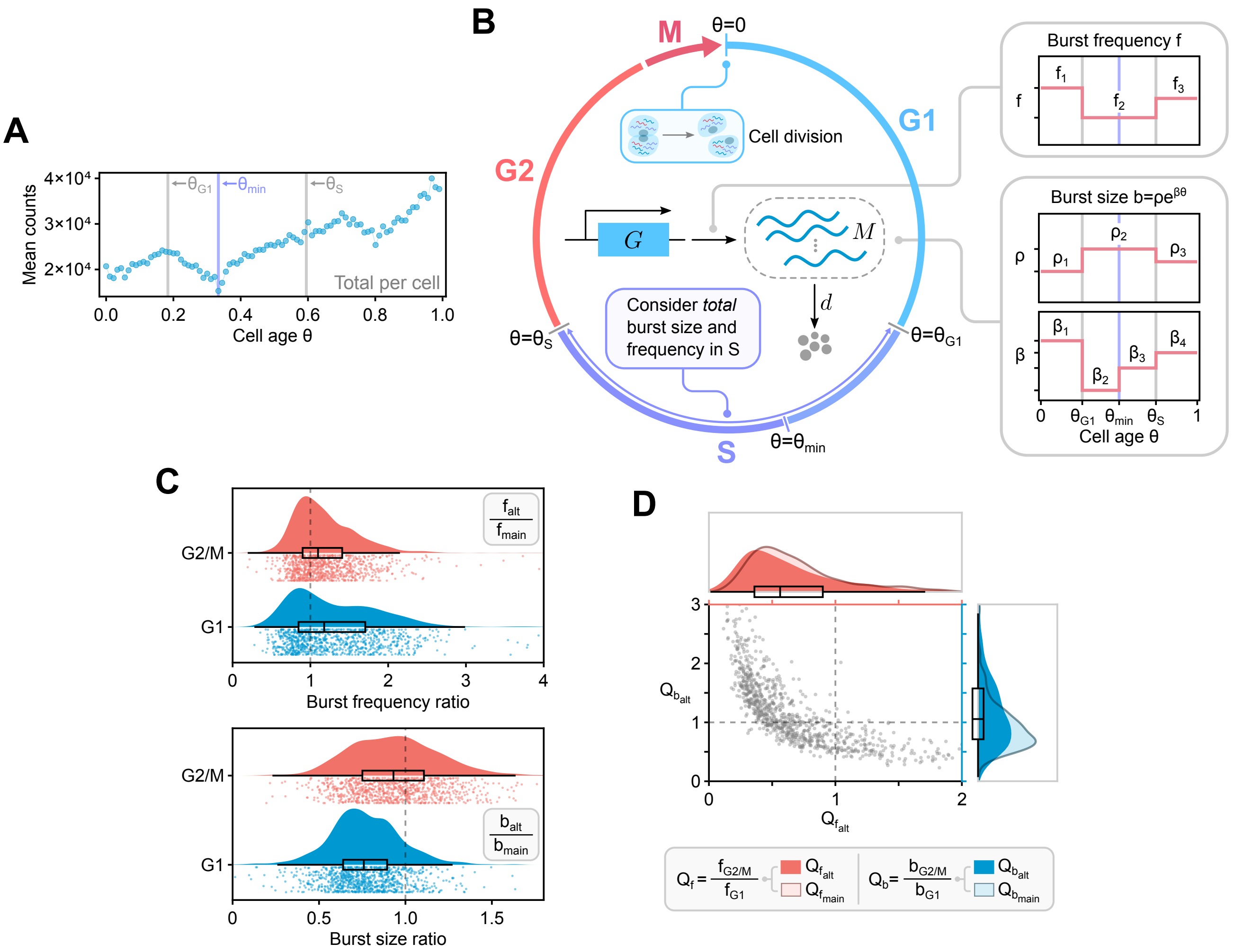


**Figure S3:** **(A)** Plot of the total mRNA counts versus cell age. The global minimum in mRNA counts during the S phase is noted by a violet line at $\theta_{\mathrm{min}}$ while the two grey vertical lines show the G1-S and S-G2/M transitions. **(B)** Schematic of an alternative age-dependent model describing bursty gene transcription with burst frequency and size that vary with cell age and cycle phase, which also takes into account the doubling of gene copy number when DNA replication occurs and mRNA dilution when cell division occurs. Note that the model presented here differs from the age-dependent model in the main text only by its piecewise parametrisation in the S phase, as discussed in Supplementary Text S3.**(C)** Raincloud plots of burst frequency (top) and burst size (bottom) ratios between the alternative and main age-dependent model estimates, inferred for G1 (blue) and G2/M (red) phases separately. Here we consider 958 genes that remain after a series of filtering steps applied to both models. **(D)** Scatter plot of the burst size ratio, $Q_{b_{\mathrm{alt}}}$, versus the burst frequency ratio, $Q_{f_{\mathrm{alt}}}$, between the G2/M and G1 phases for all 958 genes (each grey dot indicates a single gene) estimated with the alternative $\theta$-dependent bursty model. The smoothed histogram on the right (in blue) represents the distribution of burst size ratios, and the distribution of burst frequency ratios is given on the top (in red). The equivalent distributions of burst frequency and burst size ratios between G2/M and G1 obtained with the $\theta$-dependent model used in the main text (shown in Figure 4C), i.e. $Q_{f_{\mathrm{main}}}$ and $Q_{b_{\mathrm{main}}}$, are superimposed as transparent smoother histograms with darker contours for comparison.

### Supplementary Text 4: Alternative model with age-dependent burst frequency & fixed replication timing

In the construction of the age-dependent model in Methods, we discussed how the age-dependent mRNA production rate $\mu(\theta)$ (given by Eq. (10) in Methods) must be proportional to $e^{\beta\theta}$ in order to achieve a similar exponential scaling of the predicted mean transcript counts with cell age as observed in the experimental data. We enforced this constraint for the main-text age-dependent model by exponentially scaling the mean burst size with cell age $\theta$, in agreement with previous smFISH studies (see Results). However, from a modelling perspective, instead of considering the mean burst size, the same parametric constraint can be satisfied by an exponential scaling of the burst frequency with respect to $\theta$. In this section, we explore how such a change in model parametrisation affects the model predictions, comparing the main-text model with age-dependent burst size to an alternative model with age-dependent burst frequency.

The model description in the case of age-dependent burst frequency requires only a minimal alteration from the derivation of the main-text age-dependent model provided in Methods. We now model the mean burst size $b$ as a cell-cycle phase-specific constant and introduce an exponential scaling of the mean burst frequency with cell age $\theta$ so that $f = \rho e^{\beta\theta}$. Note that cell-cycle phase-specific average burst frequency is now defined as follows:

$$f_{\text{phase}} = \rho \sum_{\theta=\theta_i}^{\theta_f} g(\theta) e^{\beta\theta}, \tag{21}$$

where $\theta$ varies from $\theta_i$ to $\theta_f$ and $g(\theta)$ is the distribution of $\theta$ bins over all cells assigned to that phase (similarly to Eq. (11) in Methods).

As done previously, we construct the age-dependent model in a piecewise manner in order to account for the extrinsic noise due to the cell-cycle phase and the doubling of gene copy number upon DNA replication at $\theta = \theta_{\text{r}}$. Here we assume that the allele-specific burst parameters $b$ and $\rho$ are given by the respective constants $b_1$ and $\rho_1$ before replication, and $b_2$ and $\rho_2$ after replication. We again split the parameter $\beta$ into four parts for extra model flexibility required to fit the non-monotonic variation of the mean mRNA counts with cell age in the S phase, assuming a fixed replication time for all genes in the middle of the S phase at cell age $\theta = \theta_{\text{r}}$. A schematic illustration of the model and the piecewise changes in model parameters with cell age $\theta$ are illustrated in Figure S4A.

The time evolution of the mean mRNA counts due to all alleles in each cell-cycle phase are given by the following piecewise ODEs:

$$\partial_\theta \langle m\rangle_1(\theta) = 2b_1\rho_1 T e^{\beta_1\theta} - dT\langle m\rangle_1(\theta), \quad \theta \in [0, \theta_{\text{G1}}], \tag{22}$$

$$\partial_\theta \langle m\rangle_2(\theta) = 2b_1\rho_1 T e^{\beta_2\theta} - dT\langle m\rangle_2(\theta), \quad \theta \in [\theta_{\text{G1}}, \theta_{\text{r}}] \tag{23}$$

$$\partial_\theta \langle m\rangle_3(\theta) = 4b_2\rho_2 T e^{\beta_3\theta} - dT\langle m\rangle_3(\theta), \quad \theta \in [\theta_{\text{r}}, \theta_{\text{S}}], \tag{24}$$

$$\partial_\theta \langle m\rangle_4(\theta) = 4b_2\rho_2 T e^{\beta_4\theta} - dT\langle m\rangle_4(\theta), \quad \theta \in [\theta_{\text{S}}, 1], \tag{25}$$

where the boundary conditions are equivalent to Eqs. (17)-(20) in Methods. While the ODEs for the mean counts are identical to those of the main-text age-dependent model (except for the renamed constants), the ODEs for the variance of the mRNA counts due to all alleles in each cell-cycle phase now take a slightly different form:

$$\partial_\theta \sigma_1^2(\theta) = -2dT\sigma_1^2(\theta) + 2b_1\rho_1 T e^{\beta_1\theta}(1 + 2b_1) + dT\langle m\rangle_1(\theta), \quad \theta \in [0, \theta_{\text{G1}}], \tag{26}$$

$$\partial_\theta \sigma_2^2(\theta) = -2dT\sigma_2^2(\theta) + 2b_1\rho_1 T e^{\beta_2\theta}(1 + 2b_1) + dT\langle m\rangle_2(\theta), \quad \theta \in [\theta_{\text{G1}}, \theta_{\text{r}}], \tag{27}$$

$$\partial_\theta \sigma_3^2(\theta) = -2dT\sigma_3^2(\theta) + 4b_2\rho_2 T e^{\beta_3\theta}(1 + 2b_2) + dT\langle m\rangle_3(\theta), \quad \theta \in [\theta_{\text{r}}, \theta_{\text{S}}], \tag{28}$$

$$\partial_\theta \sigma_4^2(\theta) = -2dT\sigma_4^2(\theta) + 4b_2\rho_2 T e^{\beta_4\theta}(1 + 2b_2) + dT\langle m\rangle_4(\theta), \quad \theta \in [\theta_{\text{S}}, 1], \tag{29}$$

where the boundary conditions are given by Eqs. (25)-(28) in Methods. We solve the complete ODE system using Mathematica, and approximate the true mRNA count distribution at any cell age $\theta$ with a negative binomial parametrised by the age-dependent moment solutions, as defined in Eqs. (30)-(33) in Methods.

We fit the obtained age-dependent model to the experimental mRNA count data using the method of maximum likelihood and estimate the parameters $b_1, b_2, \rho_1, \rho_2, \beta_1, \beta_2, \beta_3, \beta_4$ for each of the 1351 bursty genes (left after initial gene filtering outlined in Methods). Note that the parameter estimation procedure and the parameter constraints are equivalent to those described for the original age-dependent model in Methods. We next discard genes with poor model fits that predict a negative correlation of the mean counts with cell age in the G1 or G2/M phases, or have a negative associated $R^2$ value for the predicted mean and/or variance throughout the entire cell cycle, as described in Methods. In addition, we discard a handful of genes that in any cell-cycle phase have an associated burst size $b$ tending to the lower parameter bound, i.e. enforcing $b_1, b_2 > 0.002$, because lower burst size values (or, similarly, very high burst frequency values) are indicative of a Poisson-like fit that can lead to physically unrealistic parameter ratio estimates. After filtering, we are left with 1087 genes, 1027 of which intersect with the 1044 genes obtained using the age-dependent model in the main text. Therefore, we consider the overlapping set of 1027 genes and their model fits for the model comparison below.

In Figure S4B, we demonstrate that the burst frequency and burst size ratios between the G2/M and G1 phases estimated using the alternative model with age-dependent burst frequency closely agree with the same burst parameter ratio estimates obtained using the main-text model with age-dependent burst size. More precisely, for the 1027 overlapping genes, the burst frequency and burst size ratios between G2/M and G1 phases predicted with the alternative age-dependent model have a median of 0.50 (1st and 3rd quartiles are 0.36 and 0.69) and 1.09 (1st and 3rd quartiles are 0.86 and 1.31), respectively, while for the main-text model the median burst frequency ratio is 0.62 (1st and 3rd quartiles are 0.43 and 0.88) and the median burst size ratio is 0.86 (1st and 3rd quartiles are 0.64 and 1.15). We also note that both age-dependent models lead to a similar negative correlation between the burst frequency and burst size ratios: for the set of 1027 genes the correlation coefficient is -0.56 using the alternative age-dependent model, and -0.51 using the main-text model. Furthermore, in Figure S4C we show for three sample genes that the predicted means, variances and distributions of mRNA counts are in good agreement with the same statistics computed from the experimental data, regardless of which of the two age-dependent models is used.

In summary, we find that the choice of an age-dependent burst frequency (and constant burst size) over an age-dependent burst size (and constant burst frequency) in our cell-age and cell-cycle phase-dependent model does not have a significant impact on the parameter estimates at the individual-gene level, and both model parametrisations lead to consistent transcriptome-wide results. Note that our preference for the model with age-dependent burst size in the main text is motivated by previous experimental findings, as discussed in Results.

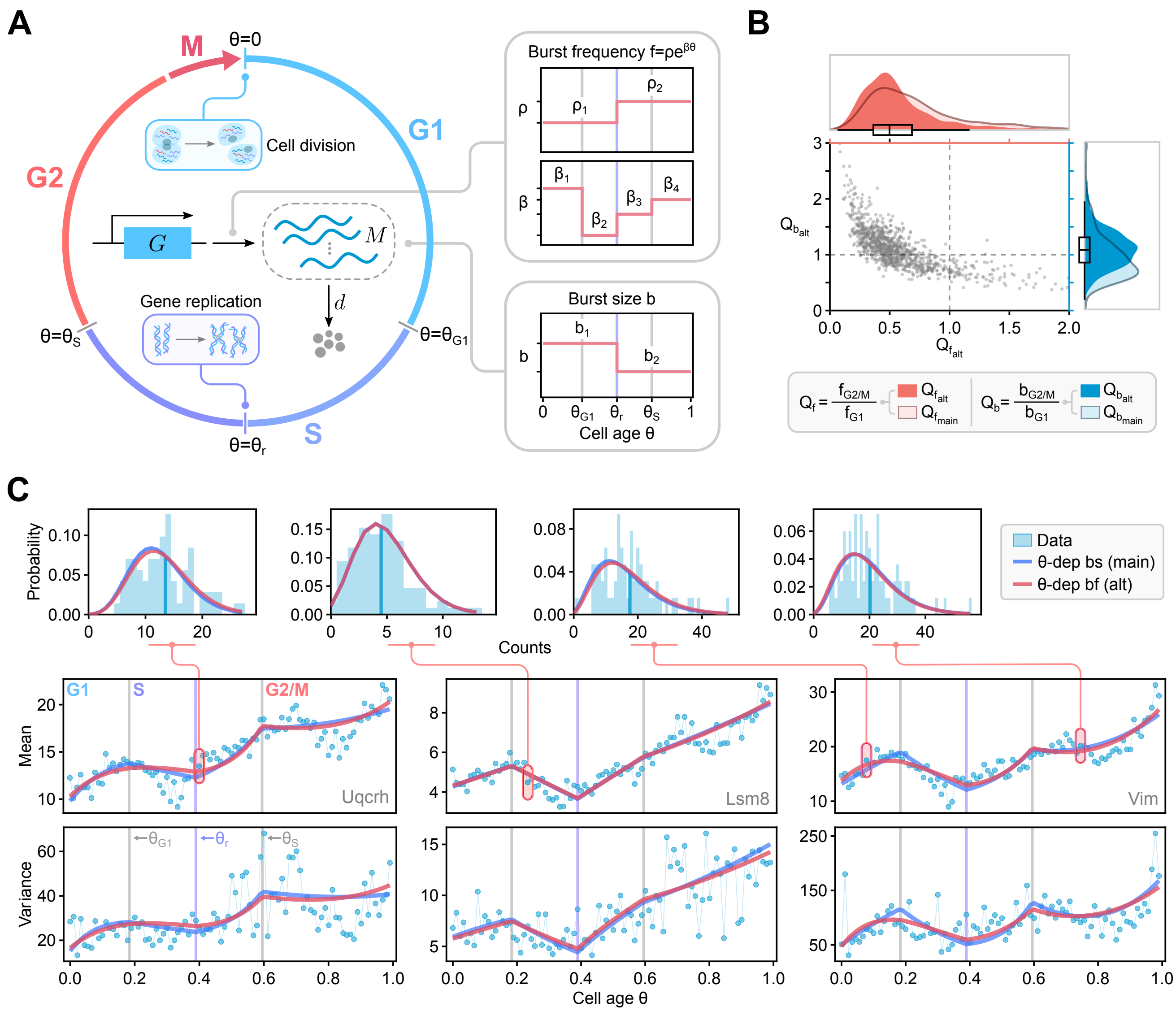


**Figure S4:** **(A)** Schematic of an alternative age-dependent model describing bursty gene transcription with burst frequency and size that vary with cell age and cycle phase, which also incorporates the doubling of gene copy number upon DNA replication and mRNA dilution during cell division. As discussed in Supplementary Text S4, the only difference from the main-text age-dependent model is the placement of the exponential scaling of transcription with cell age $\theta$: we consider age-dependent burst size in the main-text model, and age-dependent burst frequency in the alternative model. **(B)** Scatter plot of the burst size ratio, $Q_{b_{\rm alt}}$, versus the burst frequency ratio, $Q_{f_{\rm alt}}$, between the G2/M and G1 phases for all 1027 genes (each grey dot indicates a single gene) estimated with the alternative model with $\theta$-dependent burst frequency. The smoothed histogram on the right (in blue) represents the distribution of burst size ratios, and the distribution of burst frequency ratios is given on the top (in red). The equivalent distributions of burst frequency and burst size ratios between G2/M and G1 obtained with the $\theta$-dependent model used in the main text (shown in Figure 4C), i.e. $Q_{f_{\rm main}}$ and $Q_{b_{\rm main}}$, are superimposed as transparent smoother histograms with darker contours for comparison. **(C)** Comparison of the mean, variance and distributions of mRNA counts predicted by the main-text model with $\theta$-dependent burst size (blue lines, "$\theta$-dep bs (main)") and the alternative model with $\theta$-dependent burst frequency (red lines, "$\theta$-dep bf (alt)") models, with parameters inferred using maximum likelihood, and the statistics calculated from scRNA-seq data for 3 sample genes.

### References


[1] D. T. Gillespie, “Stochastic simulation of chemical kinetics,” *Annual Review of Physical Chemistry*, 58: 35–55, 2007.

[2] T. E. Loman, Y. Ma, V. Ilin, S. Gowda, N. Korsbo, N. Yewale, C. Rackauckas, and S. A. Isaacson, “Catalyst: Fast and flexible modeling of reaction networks,” *PLOS Computational Biology*, 19(10): 1–19, 2023.

[3] C. Rackauckas and Q. Nie, “DifferentialEquations.jl – A Performant and Feature-Rich Ecosystem for Solving Differential Equations in Julia,” *The Journal of Open Research Software*, 5(1): 2017.

[4] A. Waisman, F. Sevlever, M. Elías Costa, M. S. Cosentino, S. G. Miriuka, A. C. Ventura, and A. S. Guberman, “Cell cycle dynamics of mouse embryonic stem cells in the ground state and during transition to formative pluripotency,” *Scientific Reports*, 9(8051): 1–10, 2019.

[5] L. V. Sharova, A. A. Sharov, T. Nedorezov, Y. Piao, N. Shaik, and M. S. Ko, “Database for mRNA half-life of 19 977 genes obtained by DNA microarray analysis of pluripotent and differentiating mouse embryonic stem cells,” *DNA research*, 16(1): 45–58, 2009.
